# RaptGen: A variational autoencoder with profile hidden Markov model for generative aptamer discovery

**DOI:** 10.1101/2021.02.17.431338

**Authors:** Natsuki Iwano, Tatsuo Adachi, Kazuteru Aoki, Yoshikazu Nakamura, Michiaki Hamada

## Abstract

Nucleic acid aptamers are generated by an *in vitro* molecular evolution method known as systematic evolution of ligands by exponential enrichment (SELEX). A variety of candidates is limited by actual sequencing data from an experiment. Here, we developed RaptGen, which is a variational autoencoder for *in silico* aptamer generation. RaptGen exploits a profile hidden Markov model decoder to represent motif sequences effectively. We showed that RaptGen embedded simulation sequence data into low-dimension latent space dependent on motif information. We also performed sequence embedding using two independent SELEX datasets. RaptGen successfully generated aptamers from the latent space even though they were not included in high-throughput sequencing. RaptGen could also generate a truncated aptamer with a short learning model. We demonstrated that RaptGen could be applied to activity-guided aptamer generation according to Bayesian optimization. We concluded that a generative method by RaptGen and latent representation are useful for aptamer discovery. Codes are available at https://github.com/hmdlab/raptgen.

## 1 Introduction

Aptamers are short single-stranded oligonucleotides that bind to specific targets through their three-dimensional folding structure. They are analogous to antibodies and have a variety of applications, including therapeutics [1, 2], biosensors [3], and diagnostics [4]. The advantages of aptamers are that they are rapidly developed by *in vitro* generation, are low immunogenic [5], and have a wide range of binding targets, including metal ions [6], proteins [7], transcription factors [8], viruses [9], organic molecules [10], and bacteria [11]. Aptamers are generated by the systematic evolution of ligands by exponential enrichment (SELEX) [12, 13]. SELEX involves iterations of affinity-based separation and sequence amplification. This iterative process results in an enriched pool that is analyzed for candidate selection. Recent advances in high-throughput sequencing have enabled us to conduct high-throughput SELEX (HT-SELEX) to collect a vast number of aptamer candidates [14–16]. Therefore, computational approaches that efficiently process high-throughput sequencing data are critical in aptamer development.

Several computational approaches have been reported for identifying aptamers using HT-SELEX data. Aptamer identification tools utilize parameters associated with the SELEX principle, such as frequency, enrichment, and secondary structure [17–20]. Although they are useful for identifying sequences from HT-SELEX data, a variation of candidates is limited by the actual sequence existence in the data. Simulation-based methods have been reported for sequence generation [21–23]. These methods require preceding motif information and, therefore, are not suitable for identifying aptamers against an unfamiliar target. Computational approaches have also been developed to predict aptamer motifs. Motif prediction is useful not only for candidate discovery but also for aptamer development processes, such as truncations and chemical modifications. Several methods have been developed for motif detection using secondary structures [24], enrichment of subsequences during SELEX experiments [25], and emphasis on various loop regions [26]. In addition to these approaches, AptaMut utilizes mutational information from SELEX experiments [22]. As nucleotide substitutions can increase aptamer affinity, mutational information is beneficial for candidate discovery. However, while insertions and deletions are also important factors for altering aptamer activity, *in silico* methods dealing with these mutations are poorly developed. Thus, a method that generates sequences from experimental data is needed to expand exploratory space, and including motif information and nucleotide mutations confer an increased opportunity for aptamer discovery.

To develop a procedure for aptamer generation and motif finding, we focused on a neural network. As reported previously, neural networks are suitable for analyzing large datasets and are compatible with high-throughput sequencing data. DeepBind adopts a convolutional neural network (CNN) to distinguish DNA motifs from transcription factors and find sequence motifs by visualizing network parameters [27]. Recurrent neural networks can also be used for sequence discovery [28, 29]. Currently, neural network-driven generative models are being applied in a broad range of research areas. Some examples of neural network-dependent generative models include deep belief networks (DBNs) [30], variational autoencoders (VAE) [31], and generative adversarial networks (GAN) [32]. For a probabilistic generation of nucleic sequences, using long short-term memory (LSTM) was proposed to mimic sequence distribution [33]. GAN-based sequence generation methods have also been proposed [34].

In small molecule discovery, variational autoencoder (VAE)-based compound designs have been reported. VAEs learn a representation of the data by reconstructing the input data from a compressed vector [31]. Kusner *et al*. used grammar-based VAE and SMILES sequences to generate chemical structures for activity optimization [35], and Gómez-Bombarelli *et al*. utilized the representation to design chemical compounds [36]. Unlike other generative models, the VAE exploits the relationship between compressed feature space and inputs in a bi-directional manner; therefore, it is suitable for visualizing similarity-oriented classifications and emphasizing important sequence features. Utilizing the VAE to convert HT-SELEX data into low dimensional space would be useful for candidate discovery; thus, VAE-based aptamer generation systems are worth investigating. To conduct VAE modeling for HT-SELEX data, the latent space should be favorable to aptamer discovery; it captures motif subsequences, robust with substitutions, deletions, and insertions, and can easily monitor effects from the subsequences.

Here, we present RaptGen, a variational autoencoder for aptamer generation. RaptGen uses a profile hidden Markov model decoder to efficiently create latent space in which sequences form clusters based on motif structure. Using the latent representation, we generated aptamers not included in the high-throughput sequencing data. Strategies for sequence truncation and activity-guided aptamer generation are also proposed.

## 2 Materials and Methods

### 2.1 Overall Study Parameters

The VAE proposed in this study is a CNN-based encoder with skip connections and a profile HMM decoder with several training methods. Two simulation datasets containing different types of motifs were generated to assess the interpretability of the decoder. Two independent HT-SELEX datasets were subjected to the VAE, and the Gaussian Mixture Model (GMM) was used for multiple candidate selection. Furthermore, Bayesian optimization (BO) was performed based on the activities of tested sequences proposed by GMM, and sequences were truncated by shortening the model length. The process is explained in detail in the following sections. An overview is shown in Figure 1.

**Figure 1.**
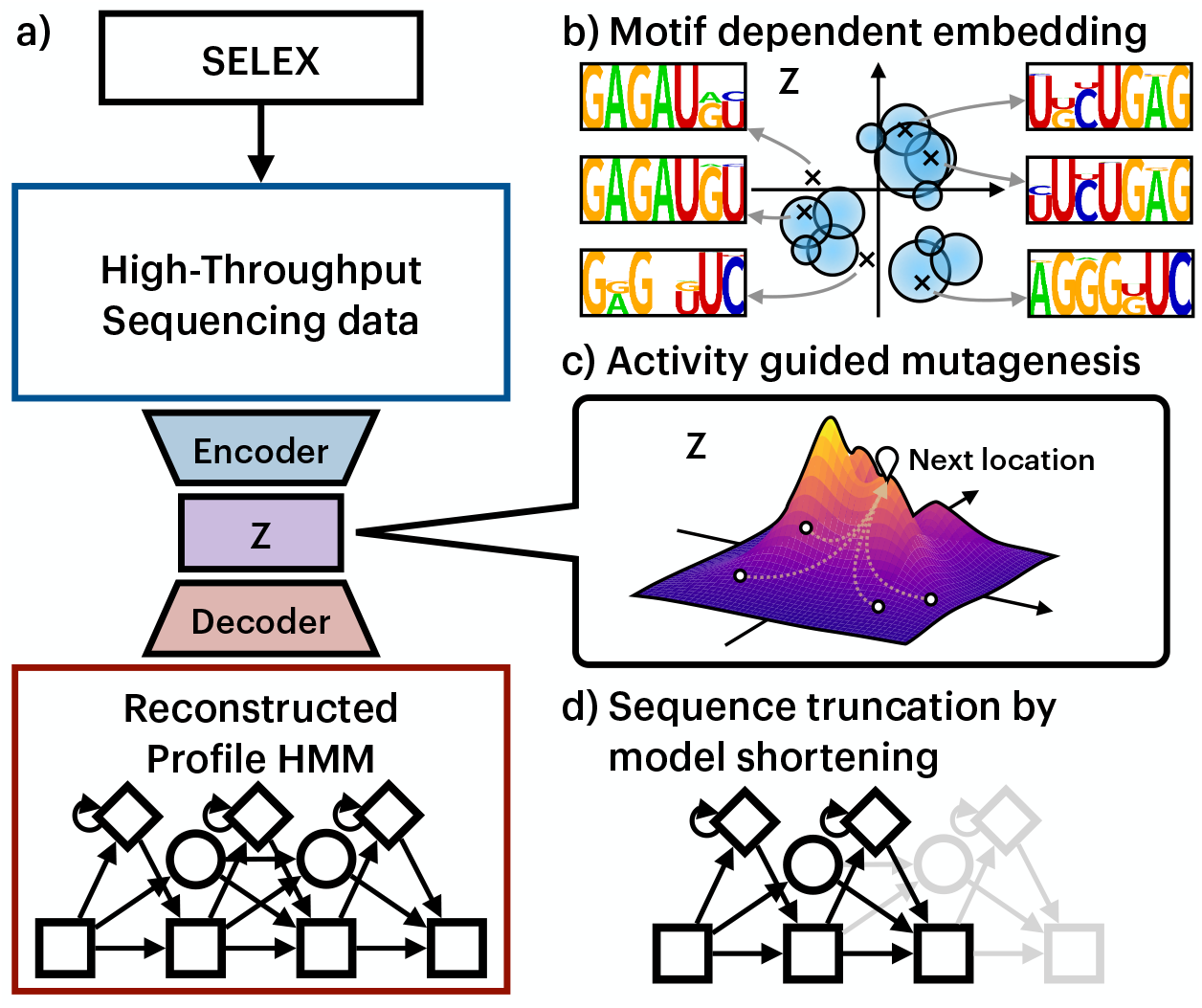
Overall RaptGen schematic and its applications. (a) RaptGen workflow. RaptGen is a VAE with profile HMM for decoder distribution, which considers insertions and deletions. Through training, RaptGen learns the relationship between high-throughput sequencing data and latent space embeddings (the latent space in shown in *Z* in this figure). (b) RaptGen can construct a latent space based on sequence similarity. It can also generate intermediate representations with no training data. (c) RaptGen can propose candidates according to the activity distribution by transforming a latent representation into a probabilistic model. (d) RaptGen can perform *in silico* sequence truncation by using a short profile HMM decoder.

### 2.2 Architecture of the RaptGen Model

#### 2.2.1 Variational Autoencoder (VAE)

VAEs consist of an encoder neural network that transforms input sequence **x** into latent distribution *q_ϕ_*(**z***|***x**) and a decoder neural network that reconstructs the input data from latent representation **z** by learning *p_θ_*(**x** | **z**). As VAE is a generative model, it can be evaluated by model evidence. However, given a dataset 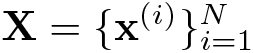, the model evidence *p_**θ**_* (**X**) is not computationally tractable. Alternatively, we can maximize the Evidence Lower BOund (ELBO); 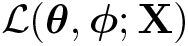 to calculate how the model describes the dataset using Jensen’s inequality,

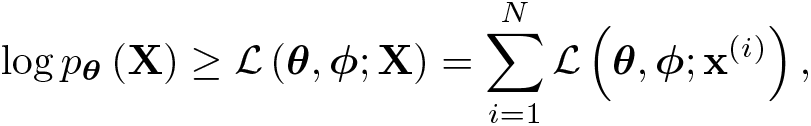

where

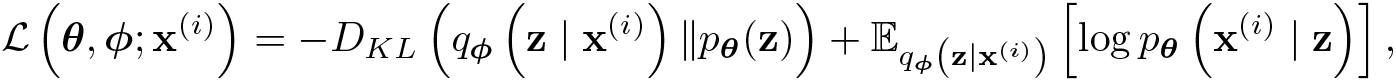

where *D_KL_*(*p || q*) is the Kullback–Leibler divergence between distributions *p* and *q*. The former term on the right-hand-side is the regularization error, and the latter is the reconstruction error. Modeling this reconstruction error to suit the problem determines the structure of the latent space.

#### 2.2.2 CNN-based Encoder With Skip Connections

The RaptGen encoder network consists of a stack of convolutional layers with skip connections. Each character was first embedded into a 32-channel vector and went through seven convolutional layers with skip connections. Then, max pooling and fully-connected layering transform the vector into the distribution parameters of latent representation *q_ϕ_*(**z***|***x**). The structure is shown in detail in the Supplementary Information subsection S1.5.

#### 2.2.3 Profile HMM Decoder Model

For modeling insertions and deletions, we used the profile hidden Markov Model (profile HMM) as the decoder for RaptGen. The profile HMM is a model that outputs by probabilistically moving from state to state (Figure 2). The profile HMM consists of a match state (M), an insertion state (I), and a deletion state (D). Each state emits specific outputs introduced to represent multiple sequence alignments [37]. The match state has a high probability of emitting a particular character, the insertion state has an equal chance, and the deletion state always emits a null character. These probabilities are called emission probabilities. The other probabilistic parameter is the transition probability. This defines the likeliness of transition from a state to the next state. In a profile HMM, the emission probability *e_S_* (*c*) is the probability of output character *c* from state *S*, and transition probability *a_S,S′_* is the probability of changing state from *S* to *S′*. These are defined as *e_S_* (*c*) = *p*(*c* | *S*) and *a_S,S′_* = *p*(*S′* | *S*), respectively.

**Figure 2.**
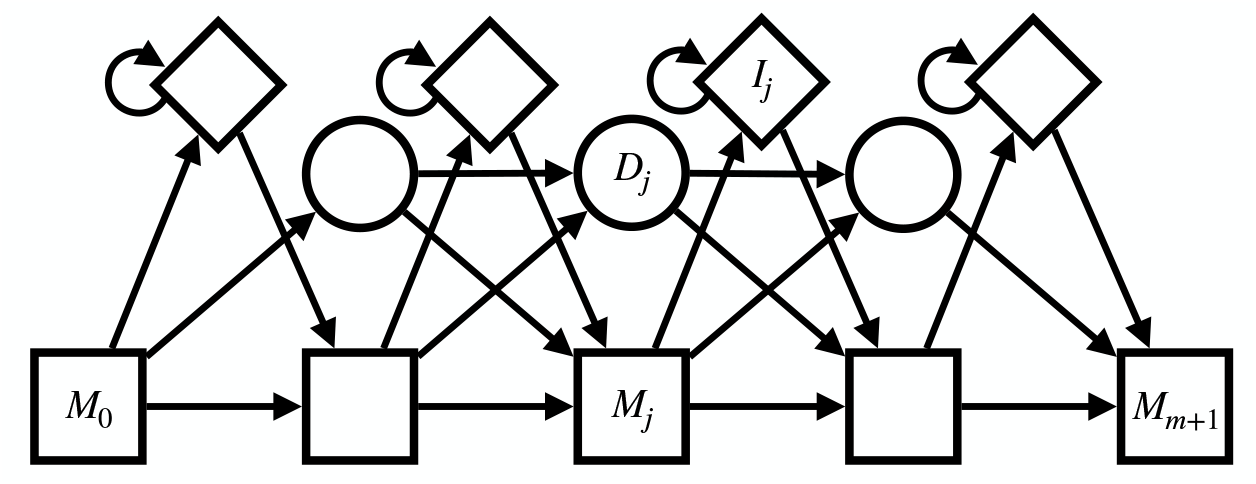
Profile HMM. The squares, circles, and diamonds represent the match *M*, deletion *D*, and insertion *I* states, respectively. The arrows represent possible transition directions of the states. The insertion and deletion states cannot go back and forth between each other. Each matching state emits a character, while the start matching state *M*_0_ and the end matching state *M*_*m*+1_ emit a null character.

Because profile HMM is a model in which the state transition depends only on the previous single state, the sequence probability *p*(**x**) can be written by using the Markov chain rule:

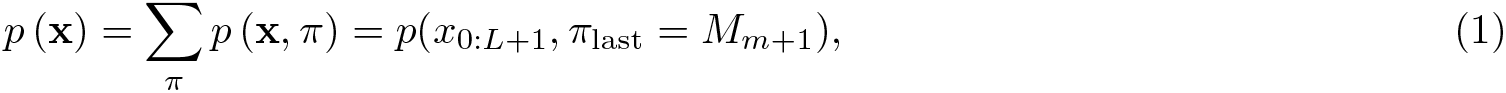

where *π* is the possible state path, *π*_last_ is the last state in the path, *L* is the length of the sequence, *x_j:k_* is the subsequence of **x** from the *j*-th character to the *k*-th character on both ends, *x*_0_ is a null character that indicates the start of the sequence, *x_L_*_+1_ is a null character that indicates the end of the sequence, and *m* is the number of matching states in the model. It is computationally expensive to calculate the sequence probability for all possible paths. Introducing a forward algorithm can lower the computational cost to 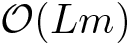. The forward algorithm consists of a forward variable defined as 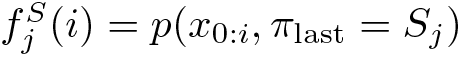, and the probability can be calculated recurrently by

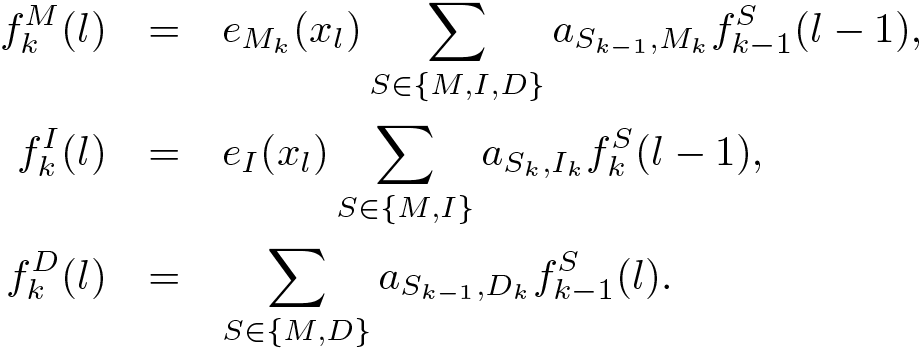

The emission probability of the insertion state does not depend on the position of the motif; therefore, it is set to a constant of 1/4 for RNA sequences. We set the probability to output the final end-of-sequence token *p*(*x_L_*_+1_*|M_m_*_+1_) to 1.

#### 2.2.4 Other Tested Decoders

Three probabilistic models were tested: the multi-categorical (MC) model, auto-regressive (AR) model, and profile HMM. The probabilistic models each have different sequence probability assignments. The MC model assigns a categorical distribution to each position of the sequence. Given the representation vector **z** and the probability of the sequence **x**, *p*(**x***|***z**) is calculated by 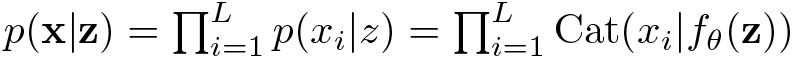, where Cat is a categorical distribution and *f_θ_* is a neural network. The AR model outputs a probability according to previous data. The probability of the sequence *p*(**x***|***z**) is calculated by 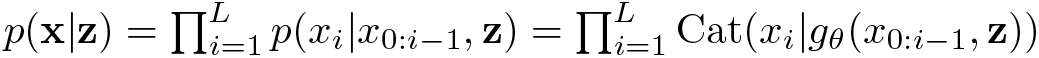, where *g_θ_* is a recurrent neural network (RNN). The architecture of networks *f_θ_* and *g_θ_* is described in the Supplementary Information subsection S1.5.

### 2.3 Training Techniques

To learn RaptGen, state transition regularization was introduced. Also, weighed regularization loss was introduced for all VAEs, including RaptGen.

#### 2.3.1 State Transition Regularization

A VAE can be trained with backpropagation by treating ELBO as a loss function. In addition to ELBO, a Dirichlet prior distribution was used on the transition probabilities to avoid unnecessary state transitions in the early rounds of training RaptGen. By penalizing transitions other than match-to-match at the beginning of the learning process, insertions and deletions are forced to occur less. This allows continuous motifs to be learned and lowers the probability of obtaining models with meaningless transitions traversing deletion states.

The probability of categorical variable ****p**** = *{p_k_}* sampled from a Dirichlet distribution is

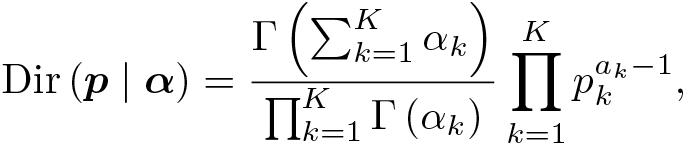

where ****α**** = {*α_k_*} is the Dirichlet distribution parameter. The regularization term is the sum of the log-odds ratio of the training probability from the matching state over each position *i*, defined as

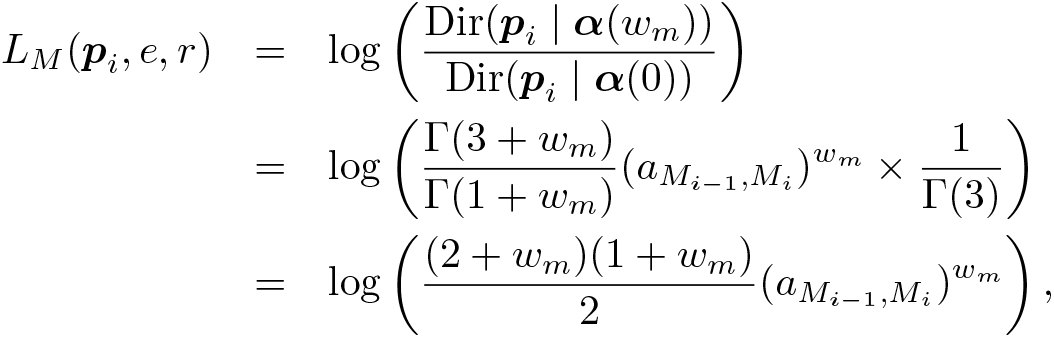

where ***p**_i_* is 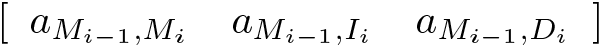 which indicates the transition probabilities from the *i*-th matching state, and ****α****(*w_m_*) = [1 + *w_m_* 1 1] is the parameter representing the induction weight *w_m_*. To make this loss zero at a specific round *R*, *w_m_* was set to 4(1 − *e/R*), where *e* is the training epoch. This regularization term was added to the ELBO during training.

#### 2.3.2 Weighted Regularization Loss

The scaling parameter for the regularization was introduced to train the VAE. Scaling the regularization term of the loss function of the VAE to minimize the value in the early epoch of training improves latent embedding [38]. The scale is defined as *e/E*, where *e* is the training epoch, and *E* is the maximum number of epochs to have scaling. After the *E* epochs of training have finished, the scale is set to 1.

#### 2.3.3 Training Settings

All sequences in the training set were filtered first. Sequences with exact matching adapters, exact matching sequence design lengths, and sequences read more than once remained. The sequences were split into training and test datasets in a 9 to 1 ratio. The model with the smallest test loss was selected through iterations. For the weighted regularization loss, the maximum number to have scaling *E* was set to 50. The state transition regularization parameter *R* was set to 50 for the profile HMM decoder. Adam was used as the training optimizer with default parameters [39]. All the networks were trained up to 2000 epochs with early stopping when the test loss was not updated for 50 epochs.

### 2.4 RaptGen Evaluation

#### 2.4.1 Simulation Data

For the simulation data shown in Figure 3(a), ten different motif sequences of length ten were generated, and single nucleotide modification with a 10% error rate was added. In other words, each motif sequence had a 10/3 percent chance of deletion, insertion, or modification at a specific position. After this procedure, sequences were randomly extended to reach 20 nt by adding nucleotides to the right and the left. We made 10,000 sequences in total, with no duplication.

**Figure 3.**
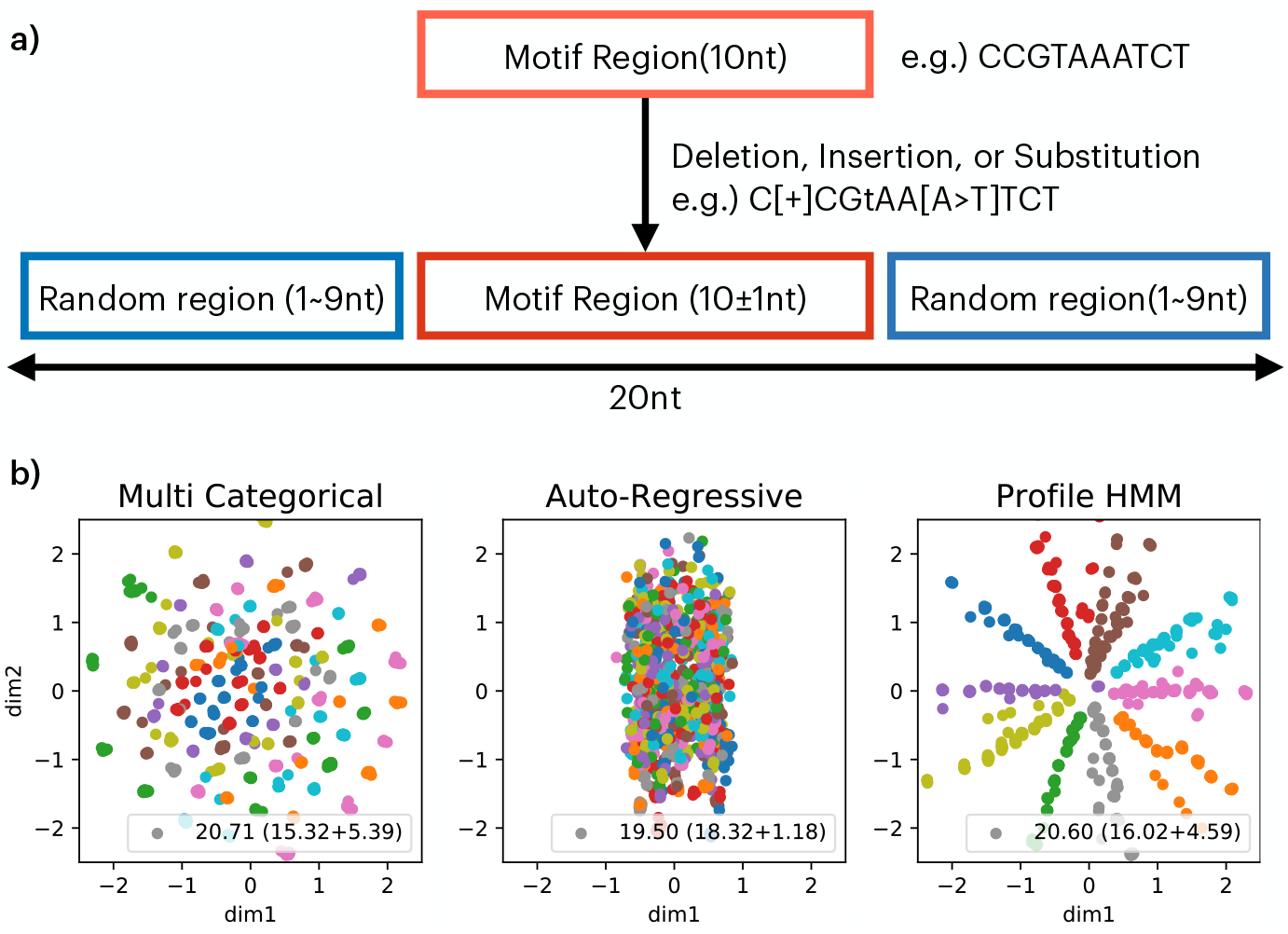
(a) Scheme of simulation data used for evaluating the decoder models. Ten different motifs with a 10% chance of having nucleotide mutations were randomly extended to 20 bases. (b) Embeddings of hypothetical motifs using different decoder models. The simulation data obtained in (a) were subjected to the VAE with the MC, AR, and profile HMM. The resulting latent space is shown. The ELBO is in the right bottom corner with the reconstructed error and KL divergence. Each motif is plotted with different colors.

For the simulation data shown in Figure 4(a), sequences containing paired motifs were generated. Two five-nt motifs were made, and then one of the motifs was randomly deleted at a probability of 25 percent each. If both motifs remained, two to six nt were randomly inserted between the left and right motifs. Subsequently, sequences were randomly extended to reach 20 nt, and 5,000 of these sequences were generated.

**Figure 4.**
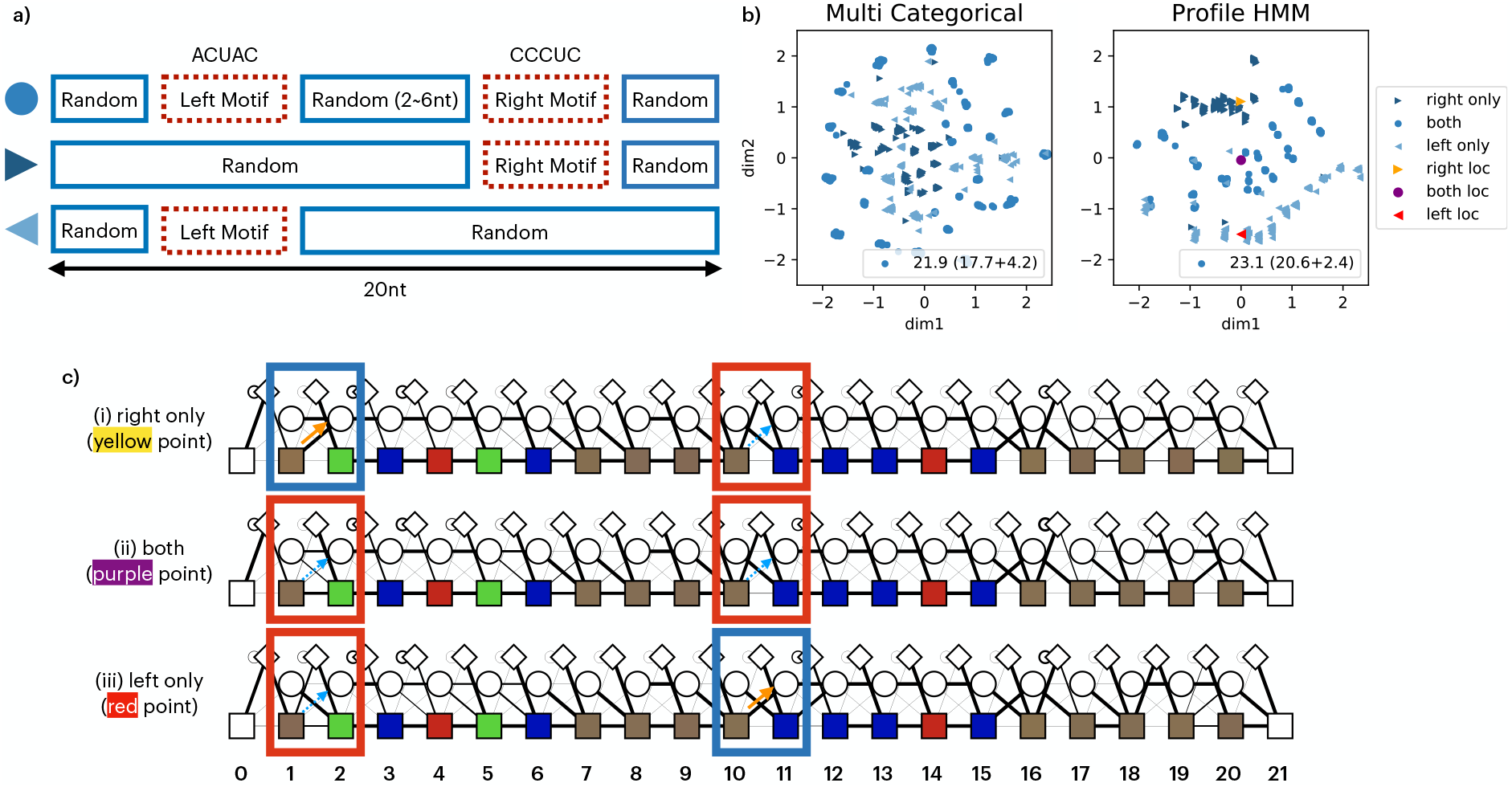
(a) Scheme for paired motifs with simulation data. A set of five nt was used with two- to six-base insertions and extended to 20 bases. Data containing one motif were also generated. (b) Embeddings of split motifs by the VAE. Simulation data generated in (a) were analyzed using the profile HMM VAE. The resulting embedding plot is shown. Plots generated by the VAE with the MC model decoder are also shown for comparison. The circle, right arrowhead, and left arrowhead represent the points from both-, right only-, and left only-motif-containing data, respectively. Yellow, purple, and red indicate representative points for each data point. The yellow and red points show the right and left motif-containing sequences, respectively. (c) Representative profile HMM obtained from the profile HMM VAE. The profile HMM indicated in (b) is shown. The thickness of the line represents the transition probability. The color of the matching state indicates the probability of the emitting nucleotide. A, U, G, and C are green, red, yellow, and blue, respectively. The blue square shows a high probability of skipping motif by moving to the deletion state, whereas the red square highlights a high probability of including the motif.

#### 2.4.2 SELEX Data

SELEX data used in this study were obtained previously [20]. The sequences are available as DRA009383 and DRA009384, which we call dataset A and dataset B, respectively. Dataset A consists of nine SELEX rounds from 0 to 8, and dataset B consists of four rounds from 3 to 6. The round with the smallest unique ratio *U* (*T*) with the restriction of *U* (*T*) *>* 0.5 was used, defined as

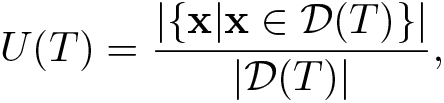

where 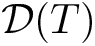 are the whole sequences, read in round *T*. The 4th round was selected for each dataset.

### 2.5 RaptGen Applications in Aptamer Discovery

#### 2.5.1 GMM for Initial Sequence Selection

We used the GMM for initial sequence selection from the obtained latent space. To efficiently select ten points to be evaluated, GMM was run 100 times with ten components, and the mean vectors of the model with the best evidence (likelihood) were selected.

#### 2.5.2 Surface Plasmon Resonance (SPR) Assay

The SPR assays were performed using a Biacore T200 instrument (GE Healthcare) as described previously with slight modifications [20]. The target proteins of datasets A and B were human recombinant transglutaminase 2 (R&D systems, Cat. No. 4376-TG) and human recombinant integrin alpha V beta 3 (R&D systems, Cat. No. 3050-AV), respectively. Aptamers were prepared with fixed primer regions and 16-mer poly(A)-tails as follows: 5’ - GGGAGCAGGAGAGAGGUCAGAUG - (variable sequence) - CCUAUGCGUGCUAGUGUGA - (polyA) - 3’ for dataset A and 5’ - GGGAGAACUUCGACCAGAAG - (variable sequence) - UAUGUGCGCAUACAUGGAUCCUC - (polyA) - 3’ for dataset B. Previously reported aptamers were used as positive controls. All evaluated sequences are listed in the Supplementary Information Table S1. Aptamers were prepared by *in vitro* transcription using a mutant T7 RNA polymerase and 2’-fluoro-pyrimidine NTPs. The running buffer consisted of 145 mM NaCl, 5.4 mM KCl, 0.8 mM MgCl2, 1.8 mM CaCl_2_, 0.05% Tween20, and 20 mm Tris–HCl (pH 7.6). A 5’-biotinylated dT16 oligomer was immobilized to both active and reference flow cells of the streptavidin sensor chip (BR100531, GE Healthcare). The poly(A)-tailed RNA was captured in the active flow cell by complementary hybridization at a concentration of 300 nm and a flow rate of 20 μLmin^*−*1^, with an association time of 60 s. The proteins were injected into the flow cells of the sensor chip at a concentration of 50 nM and a flow rate of 20 μLmin^*−*1^, with an association time of 60 s. To regenerate the sensor chip, bound aptamers were completely removed by injecting 6 m urea. Data were obtained by subtracting the reference flow cell data from the active flow cell data. The ratio of the protein-binding level to aptamer-capturing level was used as binding activity. Percent relative binding activities of positive control aptamers are shown in the results and discussion section. For normalization of dataset A, the cycle number-dependent reduction of control aptamer binding was estimated.

#### 2.5.3 Multipoint BO via Local Penalization

BO uses both the search for sequences that have not been explored to a reasonable extent and the utility of utilizing sequences with known affinity to select the next sequence for evaluation. The local penalization function is a method that can determine the multi-point expected improvement of candidates by considering the smoothness of the potential function [40]. As it converges faster than qEI [41] and other methods for simultaneous optimization. We used this method to perform multi-point optimization. Implementation was performed with the GPyOpt package [42].

## 3 Results and Discussion

### 3.1 Motif-Dependent Embeddings Using Simulation Data

#### 3.1.1 Embedding Continuous Motif with Different Decoder Models

We first attempted to construct a VAE with an encoder and a decoder applicable to aptamer discovery. In the aptamer representation space, sequences containing the same motif should be in a neighboring area. Robustness against nucleotide mutations and motif positions should also be considered. To identify a desirable decoder, we investigated different types of sequence representation models. We constructed VAEs with a CNN encoder and three different types of probabilistic models: the MC model, AR model, and profile HMM, as a decoder. Simulation data including ten different motifs were created to assess the visualizing capability of these VAEs (Figure 3a). We observed that profile HMM embedded sequences in a motif-dependent manner after training the data, whereas the MC and AR models displayed indistinctive distributions (Figure 3b). To evaluate model evidence, the ELBO was calculated. Although the MC model and profile HMM had almost the same ELBO (20.71 and 20.60) and had similar reconstitution errors (15.32 and 16.02) and KL divergence scores (5.39 and 4.59), the embedding space of the MC model failed to visualize a motif cluster. This is thought to be due to the inability of the MC model to consider motif positions. Because the nucleotide probability of each position was independently estimated in the MC model, the same motifs in the shifted position might not be aligned in latent space. The AR model had the lowest ELBO (19.50); however, the reconstitution error was the worst (18.32). Furthermore, the classification result was not optimal. We suppose that latent representation is dispensable in the AR model because the model itself has context information. Additionally, we compared the different encoder types. LSTM [43] and CNN-LSTM were evaluated in combination with the above three decoders. LSTM is used in character-level text modeling. The embedding space from the MC and AR models was still inadequate using either encoder (Supplementary Information subsection S1.8). Profile HMM created distinguishable embedding with LSTM, whereas a learning deficiency was observed in combination with CNN-LSTM (Supplementary Information subsection S1.8). Collectively, we concluded that the profile HMM decoder is favorable for motif-dependent embedding. A VAE composed of a CNN encoder and a profile HMM decoder was examined in the following study.

#### 3.1.2 Embedding Split Motifs Using the Profile HMM VAE

We next tested whether our VAE model could distinguish split motifs. Subsequence co-occurrence at distances is often observed in RNA due to intramolecular base-pairing and internal loop structures [44]. We applied simulation data with a pair of 5-nt split motifs to the VAE (Figure 4). The MC model decoder was used for comparison. Figure 4b shows the results of embedding split motifs. Plots are displayed in three groups: right motif-, left motif-, and both motif-remaining sequences. Profile HMM output sequences related to the motif, whereas the MC model scattered the sequences. We sampled representative profile HMM distributions from each population. Profile HMM visualization shows that the yellow point skips the left motif. The red point skips the right motif, both by allocating a high probability of jumping to the deletion state from the matching state (Figure 4c). Visualization of the purple point shows that the middle of two points has a low probability of skipping either of the motif fragments. The transition probability to skip the left motif 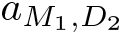 and the right motif 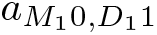 for right only-, both-, and left only-motif models were (0.995, 0.000), (0.107, 0.002) and (0.000, 0.987), respectively. Interestingly, the point located between these two motifs has a high probability of including both motifs. These results show that a profile HMM decoder is also applicable for split motifs. Hereafter, we called a VAE with a profile HMM decoder RaptGen.

### 3.2 Real Data Evaluation with RaptGen

#### 3.2.1 Examining Latent Space Dimension and Model Loss

We further evaluated RaptGen using SELEX sequence data obtained from our previous study [20]. Because real data are more complex than simulation data, we first investigated the dimensions of the latent space. Raw HT-SELEX data have 30- or 40-nt variable regions and fixed primer regions at both ends. In the present study, we used the variable region to create latent space. We tested up to 12 spatial dimensions and trained the model 50 times on dataset A (Figure 5). The minimum loss was in the four dimensions, and the second-lowest was in the two dimensions. Loss tended to increase as the embedding dimension increased; however, the loss of one-dimensional space was higher than that of the ten-dimensional space. The lower dimension would be favorable for visualization and performing BO would be advantageous, as described in later sections. Therefore, we adopted a two-dimensional space instead of a four-dimensional space for analysis.

**Figure 5.**
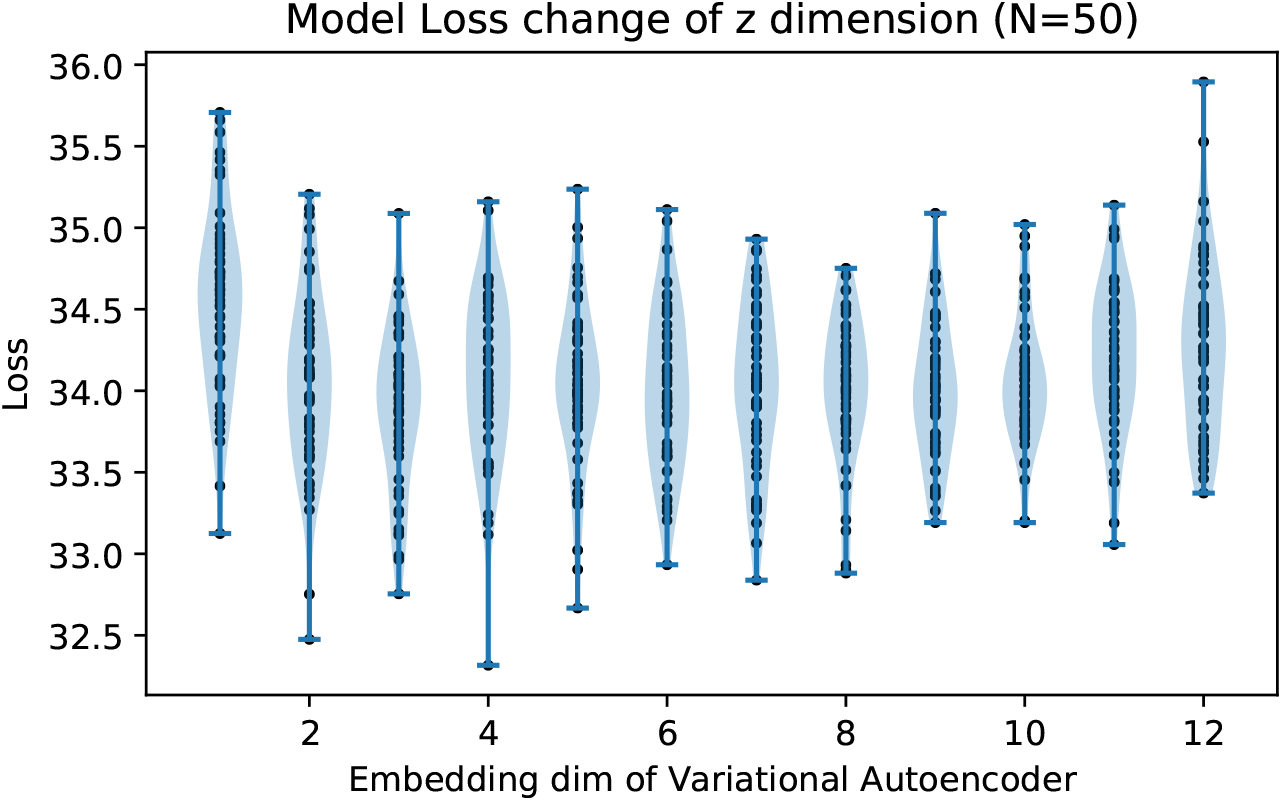
Loss over different latent dimensions. The model was trained using Data A with different dimension numbers. The minimum loss of training was plotted. Data were obtained from 50 runs with random parameter initialization.

#### 3.2.2 Embbedding High-Throughput SELEX Data with RaptGen

We next subjected two independent high-throughput SELEX datasets, Dataset A and B, to RaptGen. The resulting latent embeddings are shown in Table 1 and the Supplementary Information subsection S1.4. We previously demonstrated that aptamers from Datasets A and B exhibit continuous motifs and split motifs, respectively. As the SELEX experiment sequences are amplified with specific binding motifs, we reasoned that they would form clusters in a latent space based on their motifs. Thus, we used the GMM, which hypothesizes that data consists of a mixture of Gaussian distributions, to classify the distributions. We chose ten different points representing the latent cluster center of the GMM (Table 1). We observed that sequences with an ambiguous profile HMM such as A-GMM-2, A-GMM5, and B-GMM-0, were embedded near the latent space center. In contrast, the near edge area contained sequences that emit nucleotides preferentially. We also confirmed that similar profiles were embedded in similar areas (Supplementary Information subsection S1.4). These results provide support for the use of RaptGen to analyze high-throughput SELEX data.

**Table 1.**
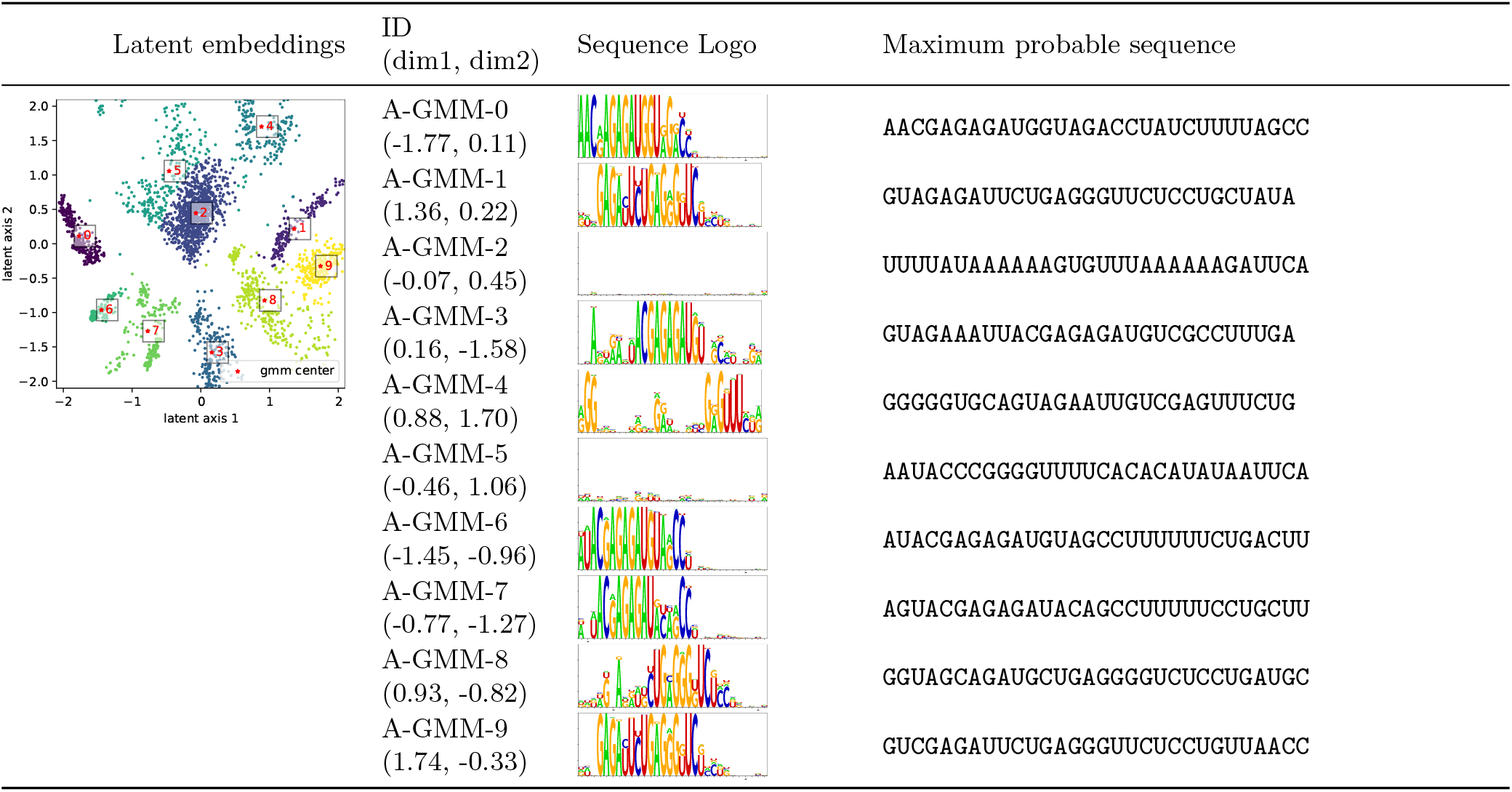
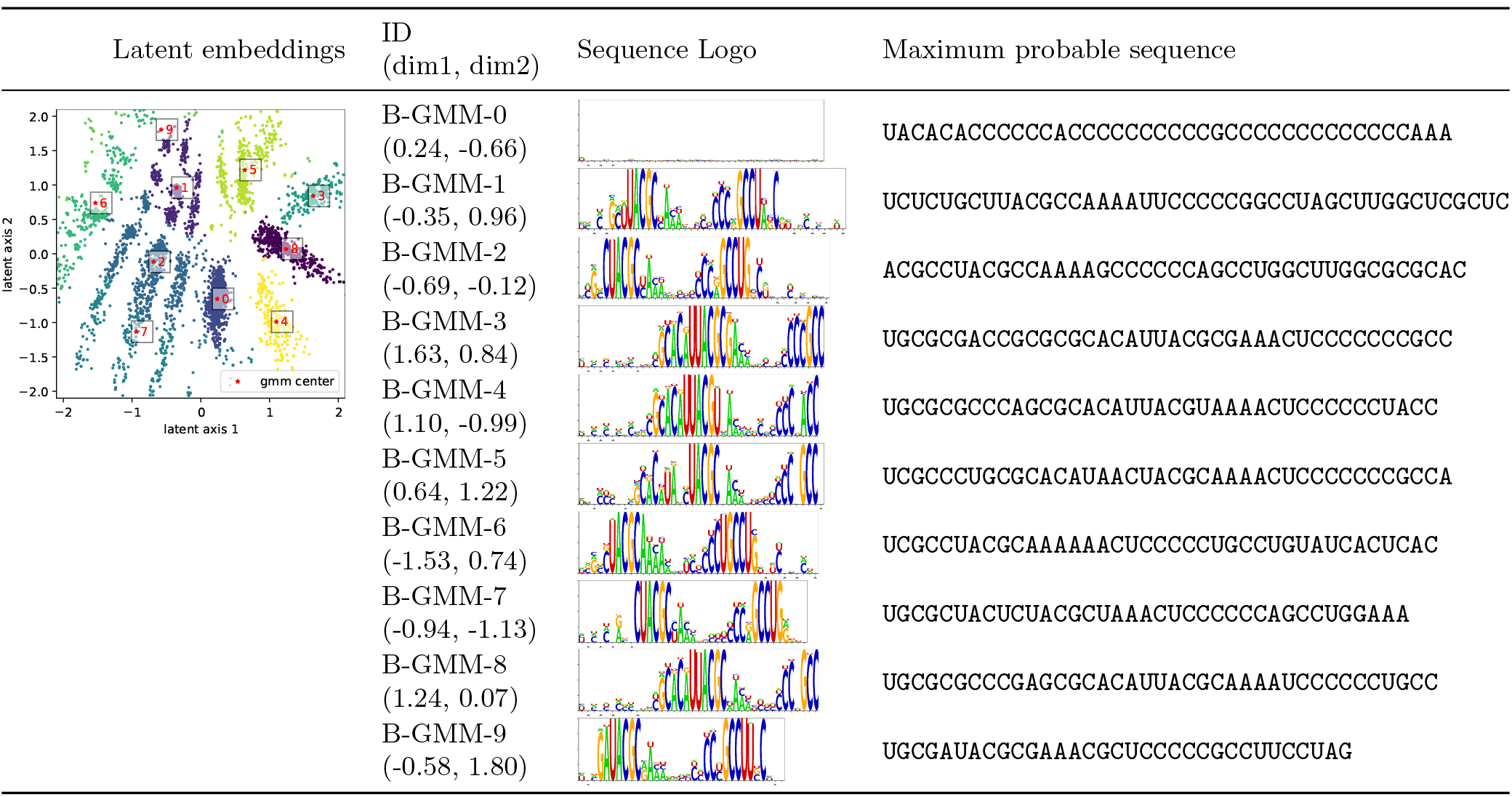
The latent embeddings and reconstituted sequences through GMM. Sequences of Data A and B were analyzed by RaptGen. The latent embeddings of Data A and B are shown in the 1st column, where clusters were estimated by GMM. The plot colors indicates the different cluster. Profile HMM were obtained from each center of the gaussian distributions. Their ID, locations in latent space, and Logo views of Profile HMM are listed in the 2nd and 3rd column, respectively. Sequences for activity evaluation were reconstituted from each Profile HMM. The maximum probable sequences were listed in the 4th column.

#### 3.2.3 Generating Aptamer Candidates from Latent Space

We attempted to generate the most probable sequence from the profile HMM of each GMM center for activity evaluation. We calculated the model state path with the highest probability and derived the most probable sequence according to the path. When the path included insertion states, we generated up to 256 sequences with no duplication by randomly replacing each insertion state with a single nucleotide and selected a sequence with the highest probability. The resulting reconstituted sequences and their probabilities are shown in Table 1. After connecting with their fixed primer sequences, aptamer RNAs were produced by *in vitro* transcription, and their binding activities were assessed by SPR. Aptamers identified in our previous study were used as positive controls. Although more than half of the candidates were found to have weak or no activity, some sequences such as A-GMM-1, B-GMM-4, and B-GMM-8 had evident binding activity. To determine whether these aptamers exist in the original data, we calculated each sequence’s edit distance from the nearest HT-SELEX sequence (Table 2). It should be noted that all candidate sequences were not included in the original SELEX data. Collectively, we concluded that RaptGen enables us to generate aptamers from the latent space and reduces the limitations of working with actual sequence data.

**Table 2.** The sequences derived by RaptGen from centers of GMM. The log probabilities of the sequences generation from Profile HMM are shown. Aptamer activities were evaluated by SPR assay. Relative activities against respective positive control are shown and the activity higher than the control are shown in bold font. The edit distance to the nearest sequence within the whole sequencing data were calculated.

| ID | Log probability | Relative activity | Edit distance |
| --- | --- | --- | --- |
| A-GMM-0 | -15.8 | 79.0 | 4 |
| A-GMM-1 | -13.3 | **107.1** | 4 |
| A-GMM-2 | -34.9 | -3.6 | 9 |
| A-GMM-3 | -16.7 | 7.0 | 3 |
| A-GMM-4 | -19.1 | 15.9 | 6 |
| A-GMM-5 | -30.1 | -0.8 | 7 |
| A-GMM-6 | -14.9 | 42.2 | 4 |
| A-GMM-7 | -17.0 | 30.9 | 5 |
| A-GMM-8 | -18.7 | 33.8 | 5 |
| A-GMM-9 | -14.5 | 74.3 | 3 |
| B-GMM-0 | -47.0 | -13.2 | 15 |
| B-GMM-1 | -35.9 | 21.9 | 11 |
| B-GMM-2 | -29.9 | -9.3 | 10 |
| B-GMM-3 | -23.9 | 74.2 | 8 |
| B-GMM-4 | -23.3 | **229.1** | 7 |
| B-GMM-5 | -26.8 | -30.5 | 9 |
| B-GMM-6 | -27.5 | -28.5 | 8 |
| B-GMM-7 | -26.4 | -7.3 | 9 |
| B-GMM-8 | -24.5 | **190.7** | 8 |
| B-GMM-9 | -23.0 | -7.3 | 8 |

### 3.3 RaptGen Application in Aptamer Discovery

#### 3.3.1 Generating Truncated Aptamers using a Short Learning Model

We proposed further applications of RaptGen for aptamer development. Since the profile HMM can handle variable sequence lengths, learning settings could diverge from the original SELEX library. For example, a decoder model does not require the same length of the random region. We attempted to generate shorter aptamers than SELEX with RaptGen. We introduced a short profile HMM with truncated length by 5 or 10 nt from the original SELEX design. Dataset A was analyzed with a 20-nt and 25-nt model (called L20 and L25), where the initial library was 30 nt. Dataset B was analyzed with a 30-nt and 35-nt model (called L30 and L35), where the initial library was 40 nt. After creating latent space, ten sequences for each length were created in a GMM-dependent manner described above. Figure 6 shows the relative activity of proposed aptamers with their lengths. For dataset A, the 28-nt candidate showed binding activity where the initial library was 30 nt. For dataset B, the 29-nt candidate showed considerable activity compared with the original setting, which was 40 nt. These results suggest that RaptGen can generate a shorter aptamer than the experimentally expected length. We found that sequences with low reconstitution probability tended to have low binding activity and that sequences showing binding activity had relatively high probability (Figure 6). This observation would be helpful for effective candidate selection. We observed a tendency of sequence extension in dataset A-L20 and L25 and dataset B-L35. For instance, in dataset A, 26 nt sequences were generated from the 20 nt RaptGen setting. We speculate that the profile HMM is prone to imitating the original length in some situations. The optimal truncation length was different for each dataset. We did not identify the cause of this difference. Further studies should be performed to determine efficient truncation.

**Figure 6.**
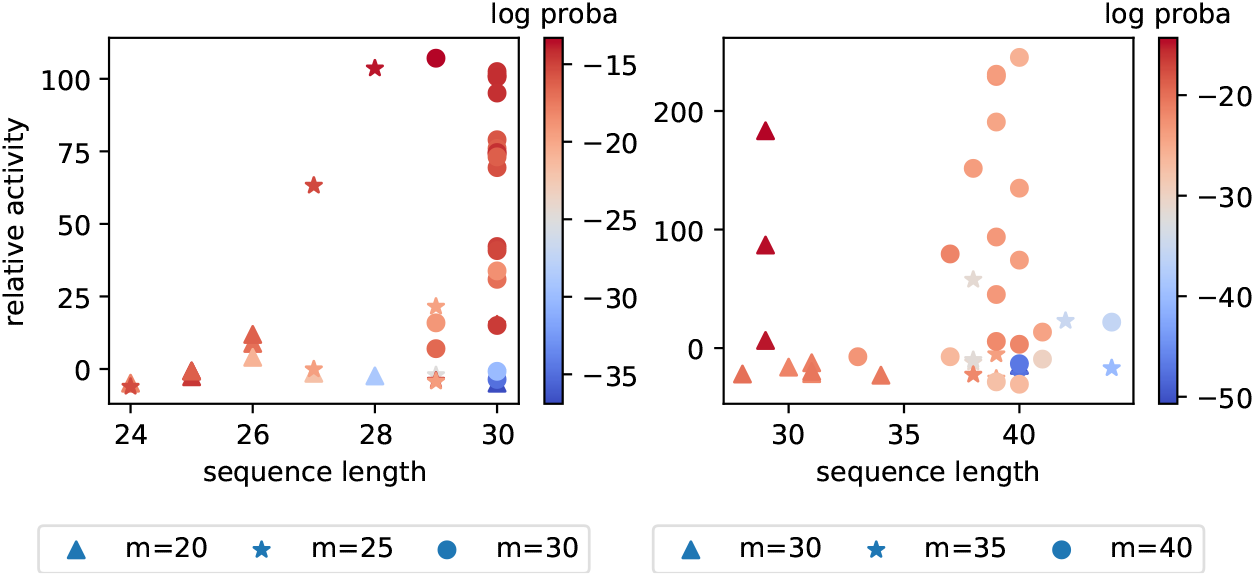
Aptamer length obtained from a short learning model. A profile HMM decoder with a length of 20 nt or 25 nt was used to analyze dataset A, in which the random region of the SELEX library is 30 nt. Similarly, a 30 nt or 35 nt decoder was used to analyze dataset B, in which the random region of the SELEX library is 40 nt. Ten candidate profile HMM were newly obtained by GMM. After reconstitution of the maximum probable sequences, aptamer activities were assessed by SPR. The scatter-plot of relative activities of aptamers and their lengths are shown, including aptamers tested in Table 1. Different markers indicate different lengths of the profile HMM decoder. Colors indicate the log probability of a sequence.

#### 3.3.2 Activity-Guided Aptamer Generation by BO

In another application of RaptGen, we generated aptamers using activity information. Aptamer derivatives harboring nucleotide mutations should be distributed around the mother sequence in the latent space. To predict effective candidates from the neighboring area of an active aptamer, binding activity distribution should be predicted. We used a BO algorithm for learning an activity distribution. Because the distribution for the BO process is required to be of low dimension, RaptGen is suitable for this strategy. To implement BO, we first embedded activity data in the latent space. The sequences listed in Table 1 were re-converted into the space. Several locations moved from the initial GMM center (Figure 7). We used these re-embedded positions to perform BO. The resulting predicted activity distributions are shown in Figure 7. To propose multiple candidates in parallel, we utilized the local penalization function [45]. Ten profile HMM were proposed and evaluated for their activity. As shown in Figure 7, candidates were generated from the peripheral area of the positive clone. We confirmed that new aptamers incorporated nucleotide substitutions (Table 3). In addition, most of them had binding activity. Similar results were obtained for both datasets A and B. These results indicate that RaptGen can propose aptamer derivatives in an activity-guided manner.

**Table 3.**
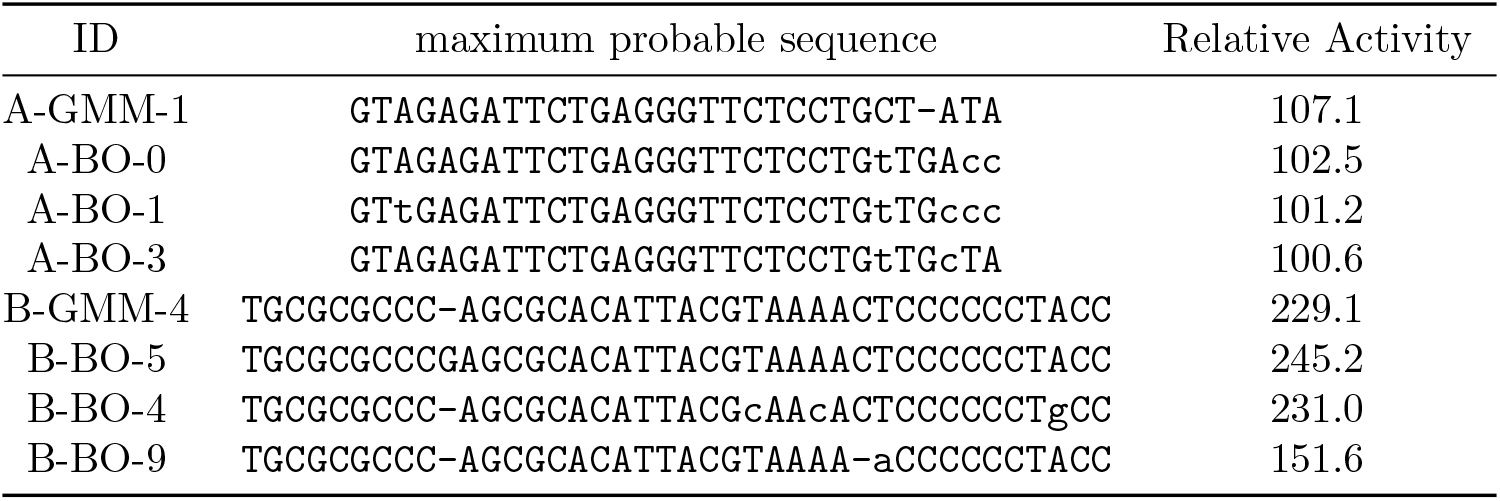
The top three sequences selected by BO and their nearest tested sequence in the embedded space. Profile HMM obtained by BO were reconstructed into a sequence by deriving the maximum probable sequence. Binding activities of the reconstructed aptamers (sequences) were evaluated by SPR experiment. Top 3 binding aptamers and their activities are shown in alignment view with nearest GMM clones. Hyphen indicates gap and lowercase letter indicates substitutions. Relative activities of A-GMM-1 and B-GMM-4 were redisplayed from Table 1 for comperison.

**Figure 7.**
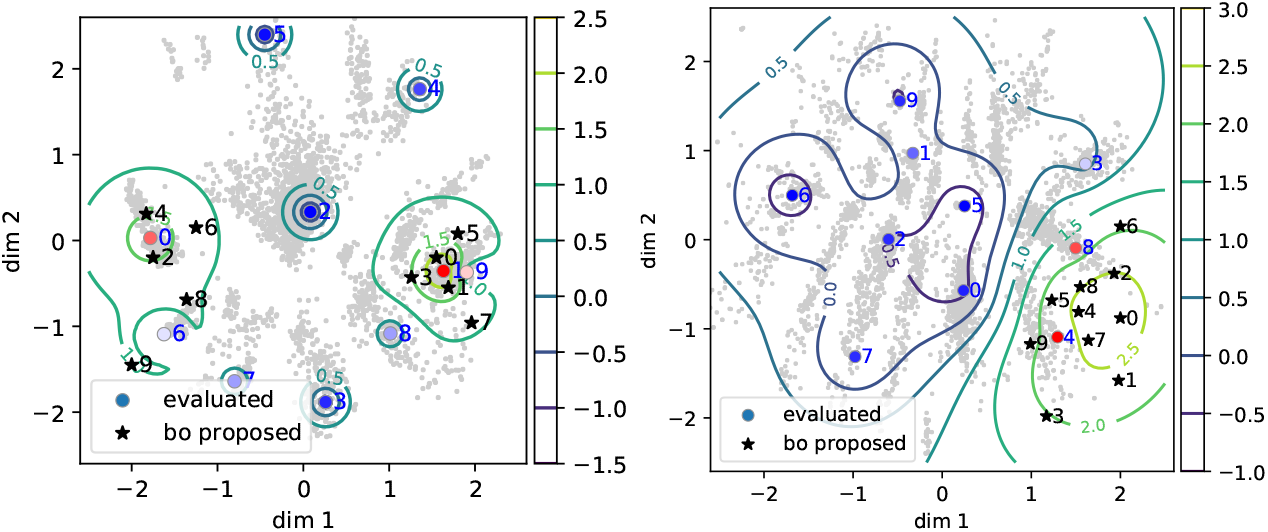
The activity distribution and proposed BO points for data A (left) and data B (right). Binding activity data shown in Table 1 were embedded into latent space. Grey points indicate latent embeddings shown in Table 1. The contour line overlaid on the embeddings indicates the predicted activity level. This is the acquisition function of BO, which is the upper confidence bound of the posterior distribution of the Gaussian process (GP-UCB) [46]. Ten points were proposed by the BO process with a local penalization function. Circles represent the re-embedded position of the GMM centers. Red and blue indicate high and low binding activity, respectively. Stars represent the locations proposed by BO.

### 3.4 Comparison with Other Techniques

Jolma and colleagues evaluated the binding specificities of transcription factors using SELEX data [8]. In addition, CNN-based research was conducted to classify randomly shuffled sequences and determine the motifs in a dataset [27]. We selected five transcription factors that were presented in these studies. We applied ten GMM distributions in this experiment and identified motifs similar to those from Jolma *et al*. Similarity was based on the edit distance of the most probable sequence from each motif. RaptGen was able to produce sequence motifs consistent with these studies (Table 4). Thus, we could search for sequence motifs obtained by running a CNN. A future study could be performed to search for an appropriate number of distributions using model selection methods such as Akaike Information Criteria (AIC) [47]. Whole GMM motifs and other results are shown in the Supplementary Information Table S3.

**Table 4.**
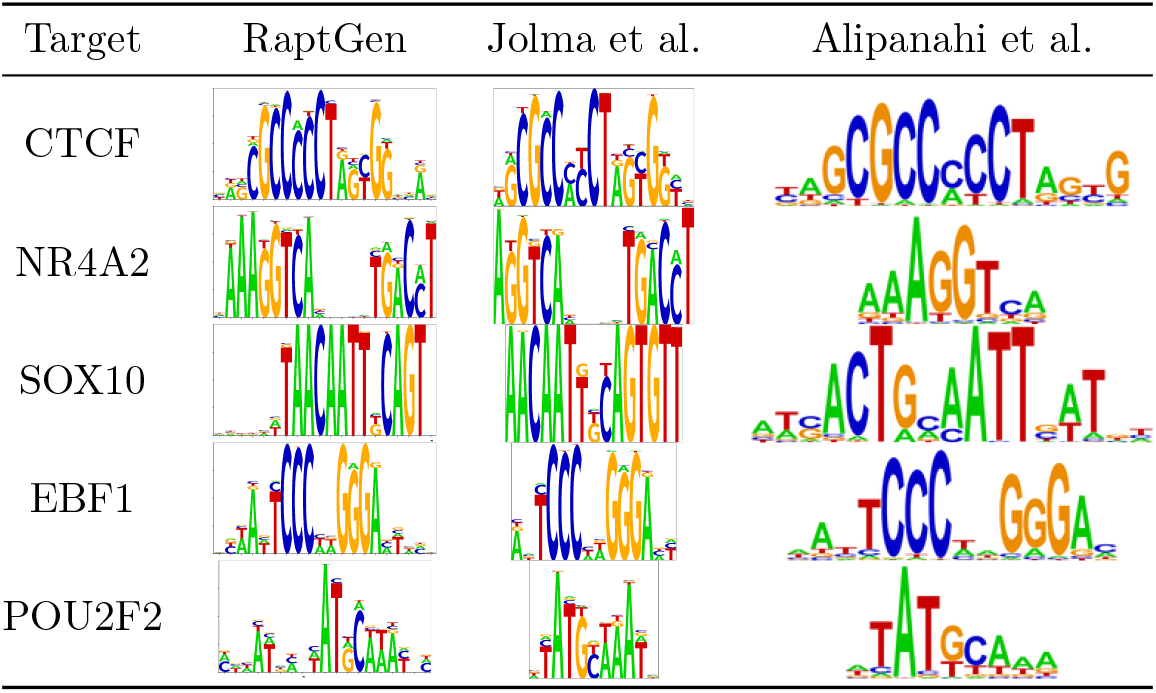
The motif similarity with previous methods. RaptGen motif was generated as described in Supplementary subsection S1.1. Motif of RaptGen is based on the emission probabilities, motifs of Jolma et al. are based on position frequency matrix (PFM) of multinomial method [15]. The motifs of Alipanahi et al. were based on the PFM, which activated specific motif scanners of the DeepBind model [27], which were derived from deepbind web service http://tools.genes.toronto.edu/deepbind/.

### 3.5 Future Work

The present version of RaptGen does not consider the secondary structure of aptamers. Secondary structure information is critical for identifying active aptamers that [19, 20]. Therefore, including the secondary structure in the sequence probabilistic model would improve RaptGen performance. An alternative model such as profile stochastic context-free grammar (SCFG) [48] will be tested in follow-up studies.

Here, we demonstrated that RaptGen could propose candidates according to activity distribution. According to BO, a sequential construction of posterior distribution would allow us to optimize activity in the latent space. For another instance of BO application, one could set the acquisition function to various indicators other than the binding activity. Therefore, we could generate candidates according to other properties of interest, including inhibitory activity against enzymes or protein-protein interactions. The application of RaptGen for this purpose is promising.

While RaptGen helps visualize and understand sequence motifs, this method requires computational cost due to sequence probability calculation. Compared with the MC model, which can calculate the sequence independently by position, and the AR model that only needs calculation on the previous nucleotides, profile HMM requires calculation on all possible state paths and previous (sub)sequences. The offset calculation cost for MC, AR, and profile HMM 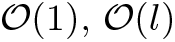, and 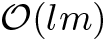, respectively, where *l* is the number of previous characters including itself, and *m* is the model length of the profile HMM. Profile HMM also needs to calculate the costly logsumexp function many times, leading to a longer training time. Additional studies are necessary to improve these issues.

## 4 Conclusion

In this study, we developed a novel nucleic acid aptamer discovery method, RaptGen, based on a variational autoencoder and profile hidden Markov model. To our knowledge, this is the first study to utilize a VAE for aptamers. RaptGen converts experimental sequence data into low dimensional space, depending on motif information. Active aptamers were generated from the representation space using a GMM. Aptamers were shorter than the original SELEX aptamers were also generated. Furthermore, we conducted a BO process for derivatizing aptamers based on activity. The creation of efficient embeddings allowed the generation of aptamers *in silico* more efficiently.

## 6 Acknowledgments

Computation for this study was performed in part on the NIG supercomputer at ROIS National Institute of Genetics. NI and MH thank members of Hamada Laboratory for their valuable comments.

## S1 Appendix

## S1.1 Method to Create Sequence Logo

To create a sequence logo for the given profile HMM, we calculated the most probable path of the profile HMM. The most probable path was iteratively acquired using the following pseudocode.

### Algorithm 1 Sequence logo generation

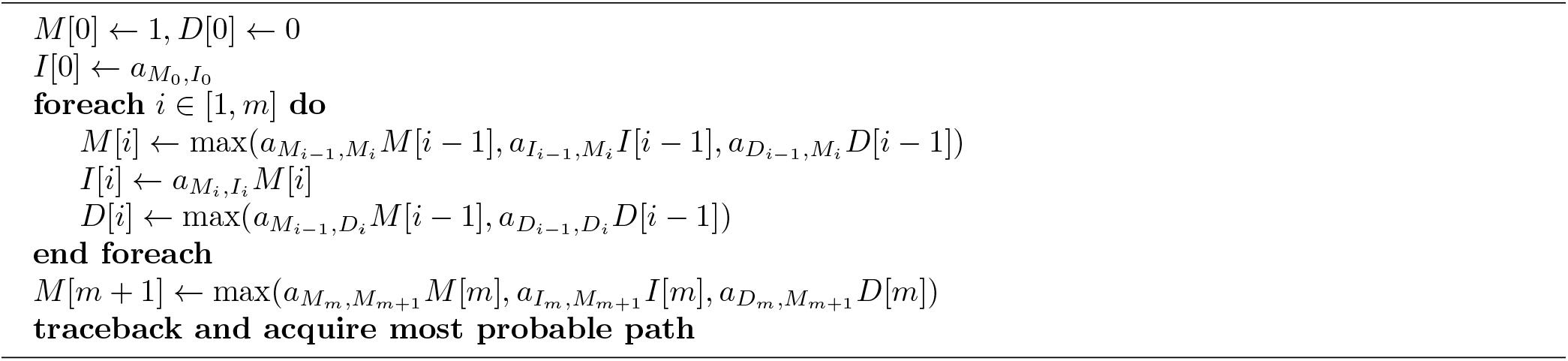

When calculating the probability of insertion state *I*, the state’s recurrency was ignored because the probability of staying on *I* recurrently is lower than that of immediately moving to the next state. After the most probable path was achieved, the sequence logo was written according to the emission probability of each state using WebLogo technology [49, 50]. The overall height *R_i_* at state *S* is defined as

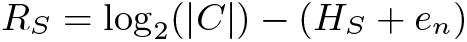

where |*C*| is the number of characters (typically 4 for RNA), *H_S_* is the uncertainty at state *S*, and *e_n_* is the correction factor. The correction factor *e_n_* adjusts the result when there are a few sample sequences. In our setting, we set *e_n_* to 0. The uncertainty *H_S_* is defined as

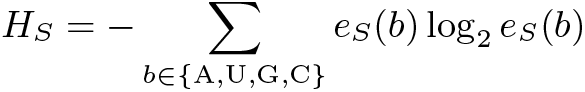

where *b* is one of the bases (A, U, G, C) and *e_S_* (*b*) is the probability of emitting base *b* from state *S*. The height of the base at a certain state *h_S_* (*b*) is defined as

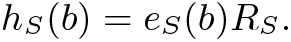

The sequence logo was written using *h_S_* (*b*), placing the higher probable base at the bottom and the state path sequence from left to right.

## S1.2 Sequence Obtained in the Present Study

The sequences obtained in the present study are shown in Table S1. The ID is named after a rule of dataID, length of the trained model, the method to select the sequence, and the index of the sequence. max seq indicates that the sequence was the most probable sequence emitted from the most probable state path of the given profile HMM. Relative activity shows % relative binding activities of positive controls assessed by the SPR experiment. The positive control sequences are as follows: A-PC is equal to Data1-11, and B-PC is equal to Data2-1 in our previous report [20].

**Table S1.**
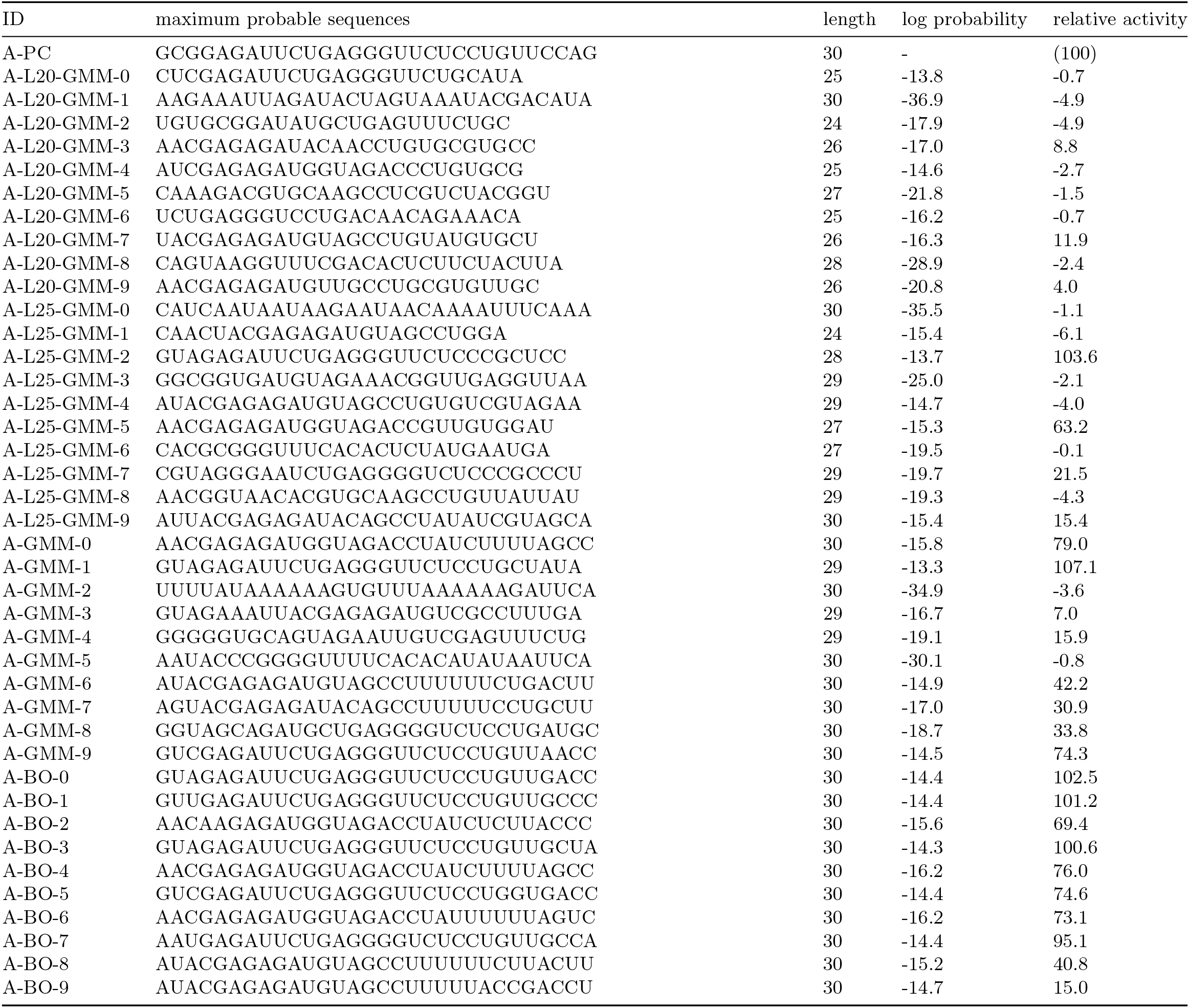
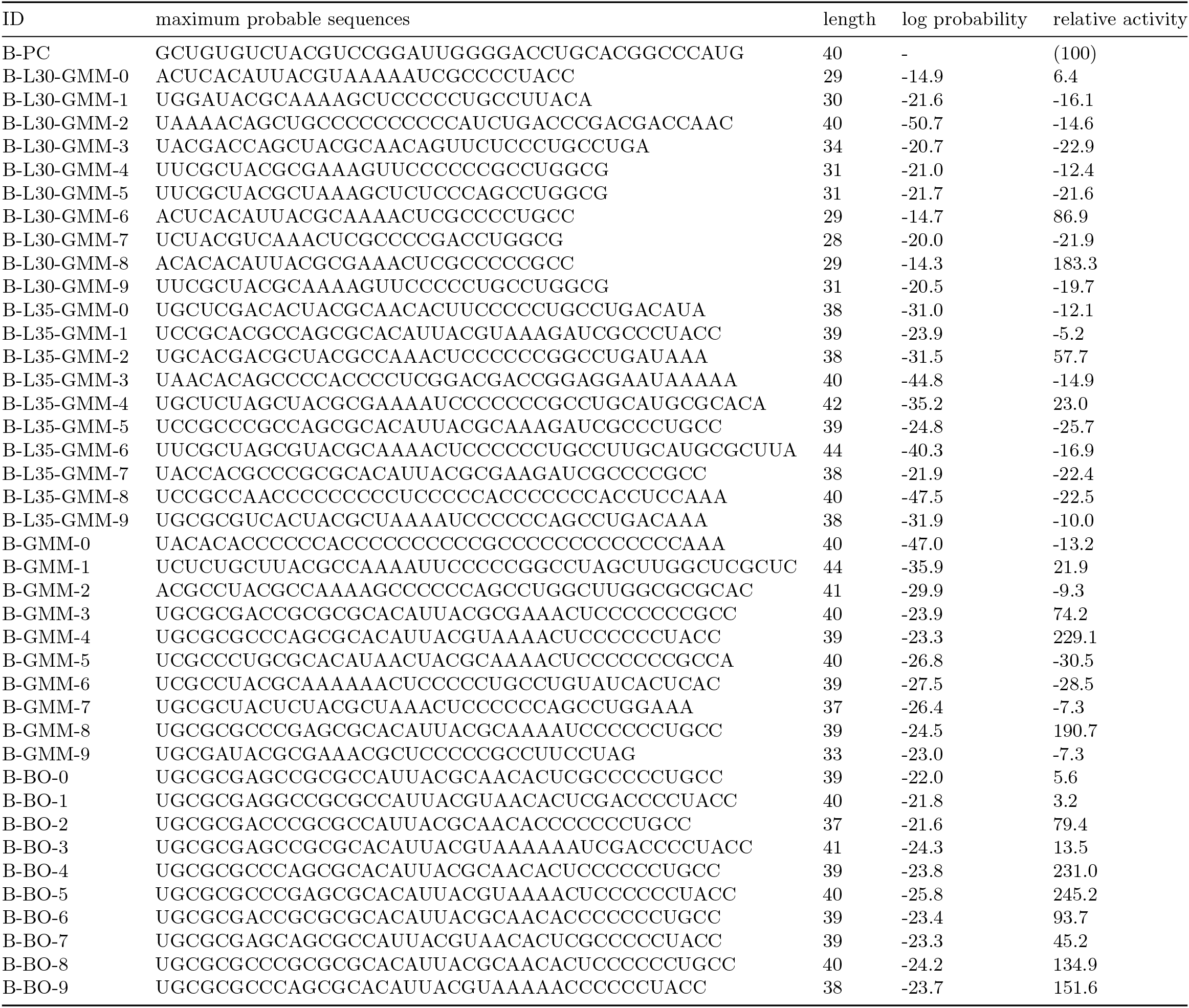
The sequences obtained through data are shown. Log probability is the probability that the sequence was obtained from a certain point of space. For GMM-, it was obtained in the initial sequence selection, and for BO-, it was obtained in the next candidate selection procedure.

## S1.3 Statistics of the Dataset

The statistics of the datasets are shown in Table S2. The column names are as follows:

- ID: The ID named after the rule; {indicator of the dataset}-{round of the SELEX}.
- raw reads: The number of reads acquired in the sequencing procedure.
- no filter unique: The number of the unique sequences with no filtration.
- adapter match: The number of sequences that match the forward- and reverse-adapter.
- designed length: The number of sequences that match the adapters and also the length of the sequence matches the experimentally designed length.
- filtered unique: The number of unique sequences that passed both adapter filtering and design length filtering.
- *>* 1: The number of filtered unique sequences that read more than once.
- *U* (*T*): The unique ratio defined in the main paper. The ratio was calculated using filter-passed sequences.
- Δ*U* (*T*): The difference in unique ratio with the previous round.

Note that the data with the lowest *U* (*T*), which holds *U* (*T*) *>* 0.5, were used.

**Table S2.** Statistics of the datasets used in our research.

| ID | raw reads | no filter unique | adapter match | designed length | filtered unique | $> 1$ | $U(T)$ | $\Delta U(T)$ |
| --- | --- | --- | --- | --- | --- | --- | --- | --- |
| A-0R | 162003 | 159606 | 114717 | 93280 | 91899 | 1353 | 0.985 | - |
| A-1R | 91610 | 90225 | 72675 | 58555 | 57595 | 929 | 0.984 | 0.002 |
| A-2R | 45431 | 44829 | 36696 | 29296 | 28856 | 424 | 0.985 | -0.001 |
| A-3R | 91441 | 90140 | 73054 | 58235 | 57349 | 857 | 0.985 | 0.000 |
| **A-4R** | 80864 | 65532 | 64503 | 51536 | 38513 | 3043 | **0.747** | 0.237 |
| A-5R | 108428 | 48575 | 86862 | 70785 | 20482 | 3760 | 0.289 | 0.458 |
| A-6R | 98237 | 25981 | 80801 | 67871 | 7750 | 2180 | 0.114 | 0.175 |
| A-7R | 49469 | 12565 | 40306 | 33101 | 3117 | 1118 | 0.094 | 0.020 |
| A-8R | 113137 | 26409 | 81177 | 67153 | 3312 | 1263 | 0.049 | 0.045 |
| B-3R | 146505 | 141174 | 102126 | 74454 | 72395 | 1874 | 0.972 | - |
| **B-4R** | 121185 | 85170 | 83310 | 58405 | 31358 | 4510 | **0.537** | 0.435 |
| B-5R | 116917 | 57404 | 83869 | 57955 | 13587 | 3375 | 0.234 | 0.302 |
| B-6R | 82488 | 37867 | 57827 | 34446 | 6064 | 1704 | 0.176 | 0.058 |

## S1.4 Sequence Logo Map

We created a map of sequence logos for the two sets of data acquired using the sequence logo creation method, as mentioned in subsection S1.1. This sequence logo is a visualization of the profile HMM at a point equally divided from −2.5 to 2.5 on each axis of the two-dimensional latent space. The sequence logo map for data A is shown in Figure S1, and for data B is shown in Figure S2.

**Figure S1.**
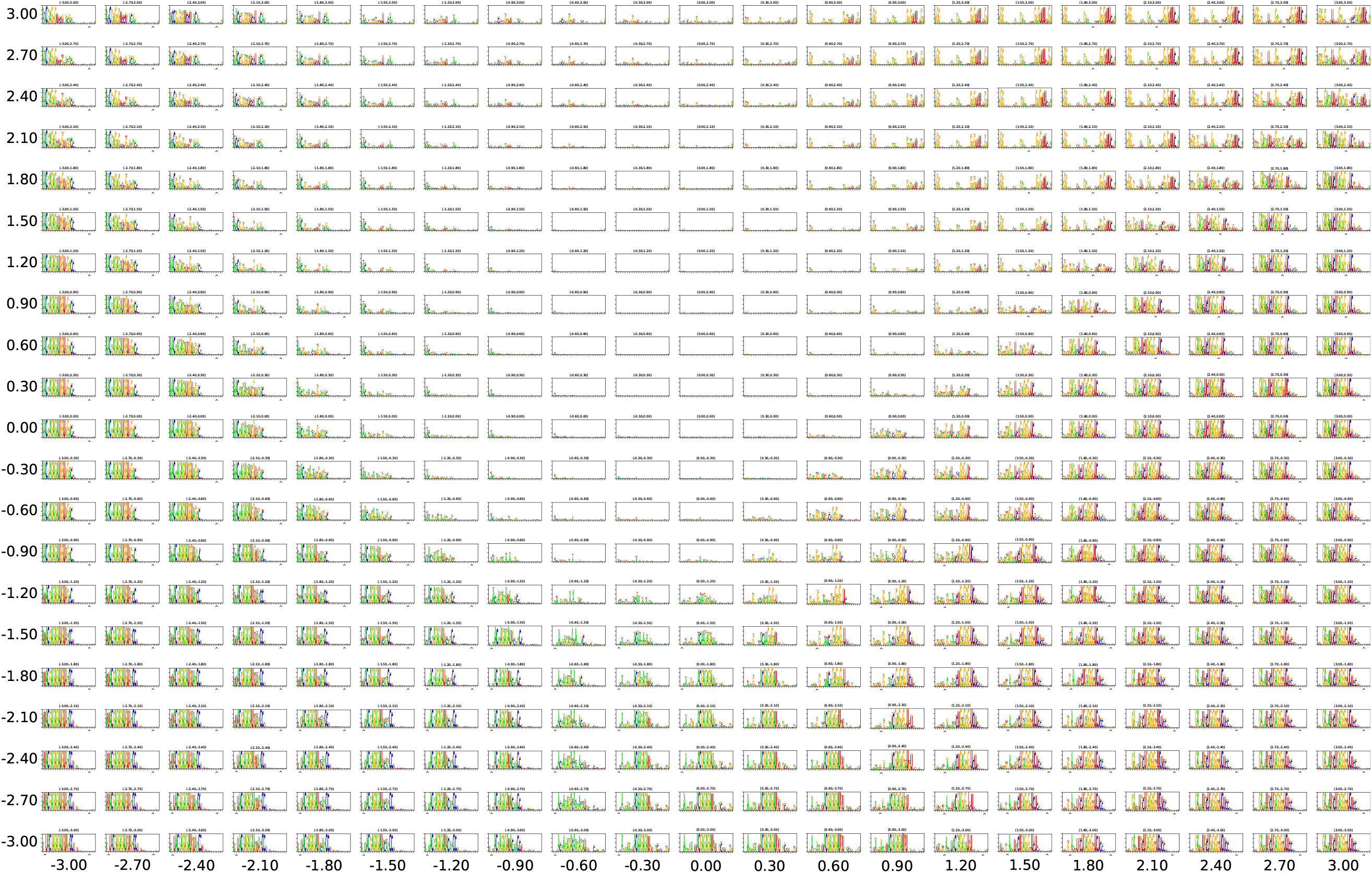
The sequence logo map for data A. The continuous motif indicated that the sequences in the data had the preference for specific subsequences. The result was partly consistent with our previous research [20] that implied AGAGAUGGUA was the truncated motif.

**Figure S2.**
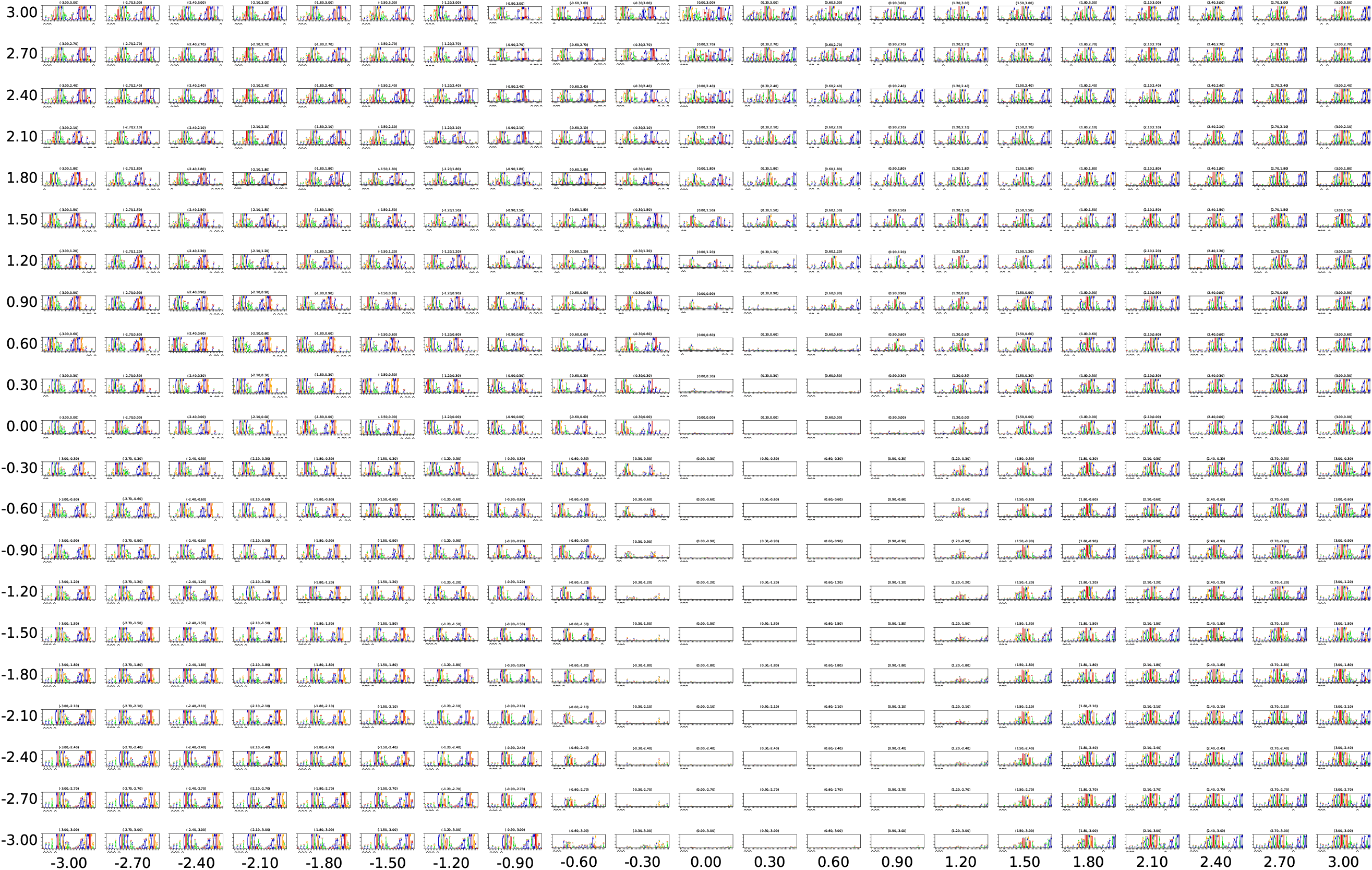
The sequence logo map for data B. The logos indicate that the Profile HMM at the center of the latent space had no preference to emit sequences while the outer surrounding points had captured split motifs.

## S1.5 Structure of RaptGen

The structure of RaptGen is shown in Figure S3. The RaptGen consists of a convolutional neural network (CNN)-based encoder and a profile HMM for decoder distribution. We further tried recurrent neural networks with long short-term memory (LSTM) and CNN-LSTM, a network of CNN followed by LSTM. The CNN utilized skip-connection [51], which enables deeper layers to learn appropriately. The models implemented in this study are available at https://github.com/hmdlab/raptgen.

## S1.6 Encoders

## S1.6.1 Convolutional neural network

A CNN captures sequential motifs that are aligned in certain positions. The encoder CNN is a network that first embeds each character into a 32-dimensional vector. Then, the six layers of the skip-connection layer capture the interactions. Finally, the max-pooling layer outputs the resulting feature vector. The skip-connection layer is a combination of a convolutional layer, batch normalization [52], and leaky rectified linear unit (leaky ReLU) [53]. The structure of the skip-connection layer is

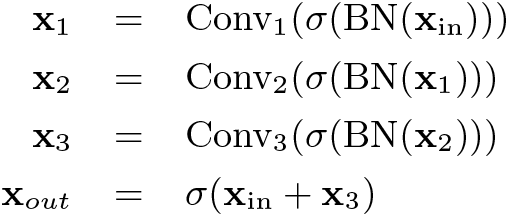

where **x**_in_ is the input vector, BN(·) is the batch normalization layer, and *σ*(·) is the leaky ReLU layer. Convolutional layer Conv_1_, Conv_2_, and Conv_3_ transforms vector dimensions from 32 to 64, 64 to 64, and 64 to 32, respectively. All convolutional layers had the same kernel size of 7 and a zero-padding length of 3 on both edges. Finally, max-pooling, taking the maximum value along the sequence, was performed, resulting in a 32-dimensional feature vector. An encoder CNN was used in the RaptGen architecture.

## S1.6.2 Recurrent neural network with long short-term memory

A recurrent neural network can consider the context of the input sequence. LSTM is an artifact that can handle the gradient vanishing problem, the hardness to learn long sequential data [43]. We used bidirectional LSTM for encoding sequences. The sequence was first embedded into a 32-dimensional vector, similar to the CNN encoder, and then it was calculated through a 16-dimensional hidden vector in each direction. The final hidden vector for each direction was concatenated into a dimensional vector, and this was used for the feature vector.

## S1.6.3 CNN-LSTM

The convolutional layer and recurrent layer are used in combination to consider both fixed-length and long-distance interactions. The CNN was almost the same as described in the previous section, with the difference that the final max-pooling was removed. LSTM treated this feature vector as sequence embedding as described in subsubsection S1.6.2.

## S1.7 Decoders

## S1.7.1 Multi categorical model

The multi-categorical model gives the probability to each position of the fixed-length sequence. The output of the model is

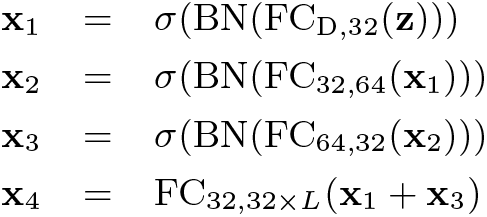

Where **z** is the sampled vector and FC_*J,K*_ is a fully connected layer, which is defined as 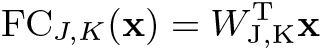 with the learnable parameter 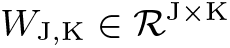. To interpret the interactions of each other, the embedding parameter is calculated using the transposed convolution function [54], which is generally the opposite of a convolutional function. The final output **x**_out_ is

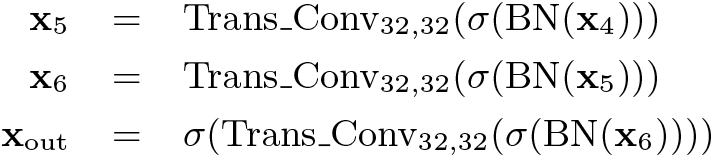

where Trans Conv_J_,_K_(·) is the transposed convolutional function with the trainable parameters of the input dimension *J* and output dimension *K*. All transposed convolutional layers have the same kernel size of 7 and the same padding length of 3.

## S1.7.2 Autoregressive model

To run the autoregressive model, we used a gated recurrent unit (GRU), which is a simplified version of LSTM [55]. The GRU is calculated as follows:

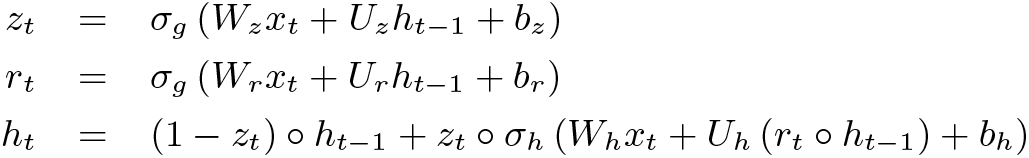

where *t* is time, *x_t_* is the input vector, *h_t_* is the output vector, *z_t_* is the update gate vector, *r_t_* is the initializing gate vector, and *W*, *U*, and *b* are parameter vectors. The probability of output sequence 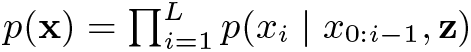 is calculated by

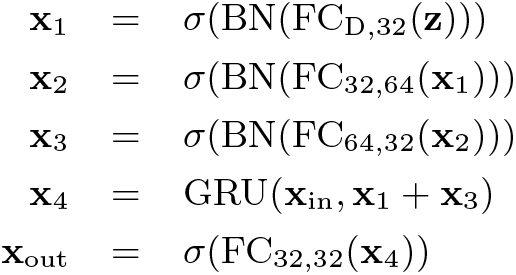

where GRU(**x***, h*_0_) is defined as a GRU function with input vector **x** of length *L* and initial hidden vector *h*_0_, which outputs *h_L_*.

## S1.7.3 Profile hidden Markov model

The profile HMM is described in Figure S3b. The embedded sequence is a *D*-dimensional vector, which is transformed into 32 dimensions by a fully connected (FC) layer. After the vector is rectified by leaky ReLU, the vector is transformed into a certain shape to fit the parameters of Profile HMM. For each parameter, FC, leaky ReLU, FC, and reshape procedure is performed.

**Figure S3.**
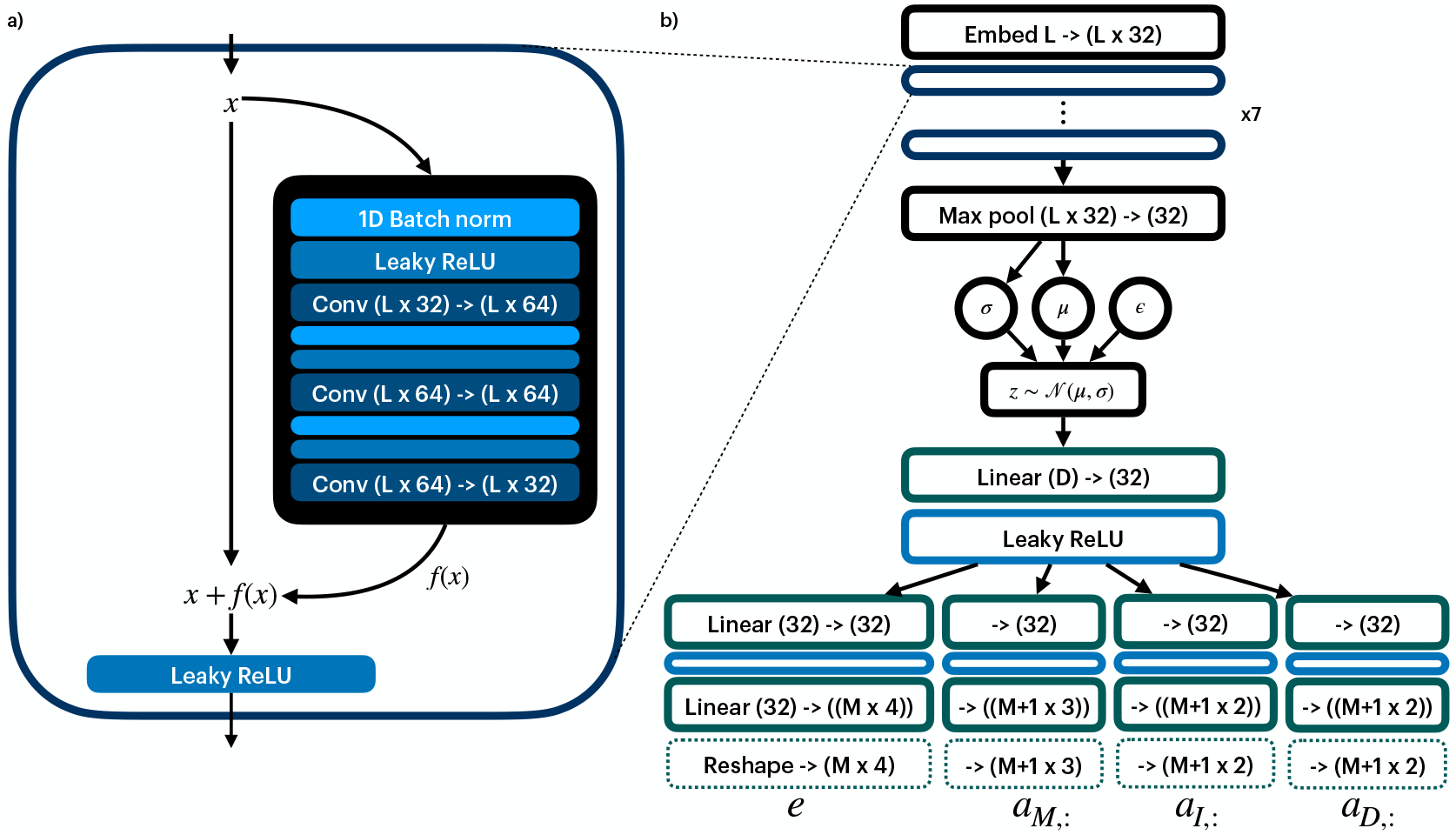
(a) The skip-connection layer. The input feature vector is first passed through 64 hidden layers, then through 32 layers, and added to the original vector. It then passes through the Leaky ReLU normalization layer and produces output. (b) Overall architecture of RaptGen. The input sequence is initially embedded into 32 feature vector and goes through skip-connection layers. After the latent mean and log-variance are calculated with the fully connected layer, which is written as “Linear,” the latent variable is sampled and calculated to fit the profile HMM parameter shapes.

## S1.8 The Embeddings of Different Encoder and Decoder Combinations

The embeddings are shown in Figure S4. The embedding of the multi-categorical probabilistic model tends to place sequences near the same motif; however, the nearest surrounding sequence is not from the same motifs. Although the autoregressive model has a lower loss, it tends to have an unsplit latent space.

## S1.9 Comparison with Other Experiments

In 2013, Jolma et al. conducted a large-scale experiment to identify transcription factor binding sites using SELEX experiments [8]. We utilized the data from that research and estimated the latent embedding, and checked whether the derived motif logo was consistent. We selected five targets whose logo was mentioned in the research of DeepBind [27]. Table S3 shows learned embedding spaces and the sequence logo of the GMM center trained on 10 components. Although the embeddings did not clearly split into 10 areas, the Profile HMM logo was consistent with previously determined motifs. Logo images of previous research were downloaded from the CisBP database [56]. The motif learned by Deepbind with the top three weights is also shown.

**Figure S4.**
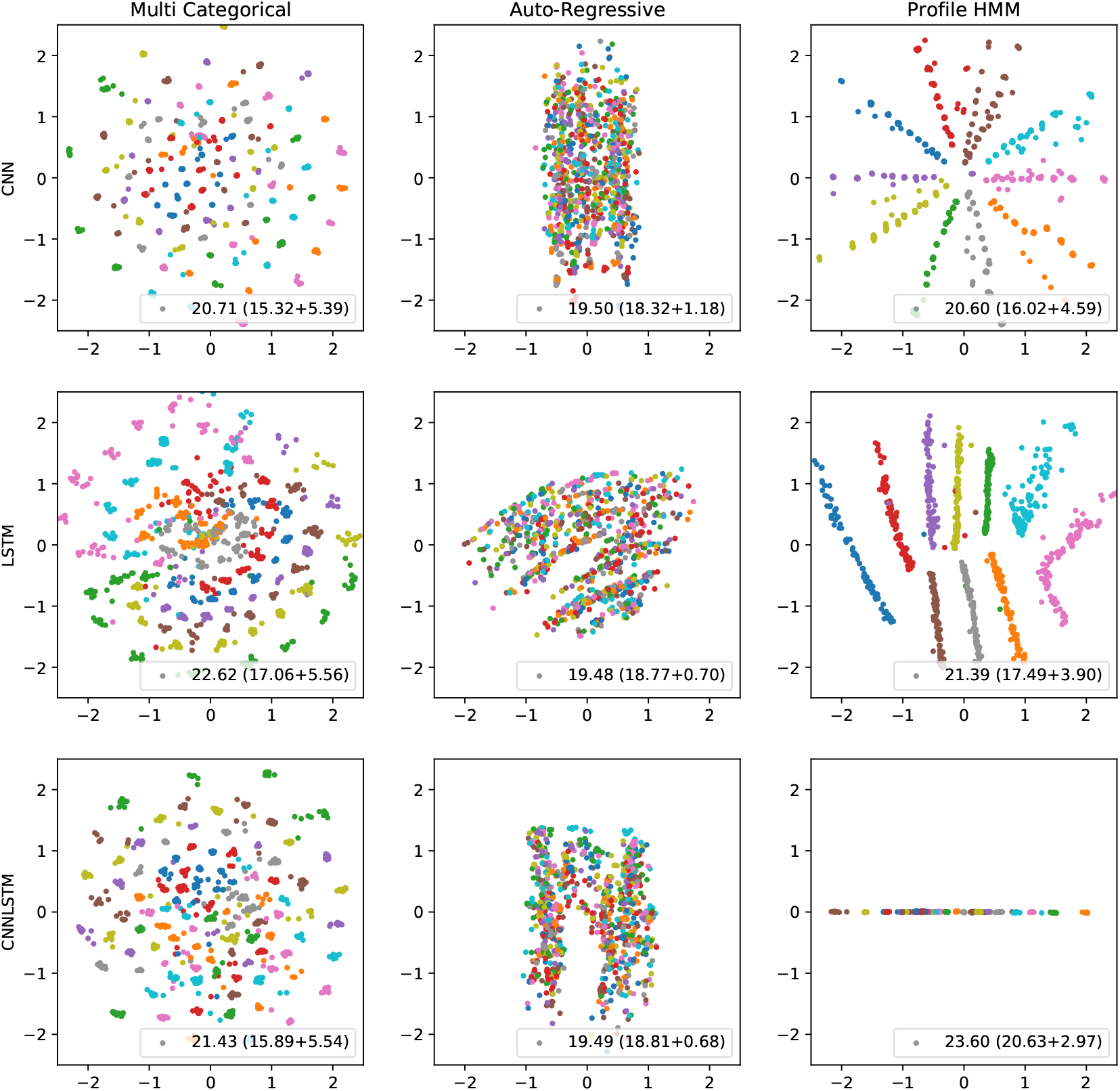
Embeddings of different encoders and decoders. The minimum loss is written at the bottom-right corner. The reconstruction error is to the left, and the regularization error is to the right in the braces. The color represents the motif that each sequence contains.

**Table S3.**
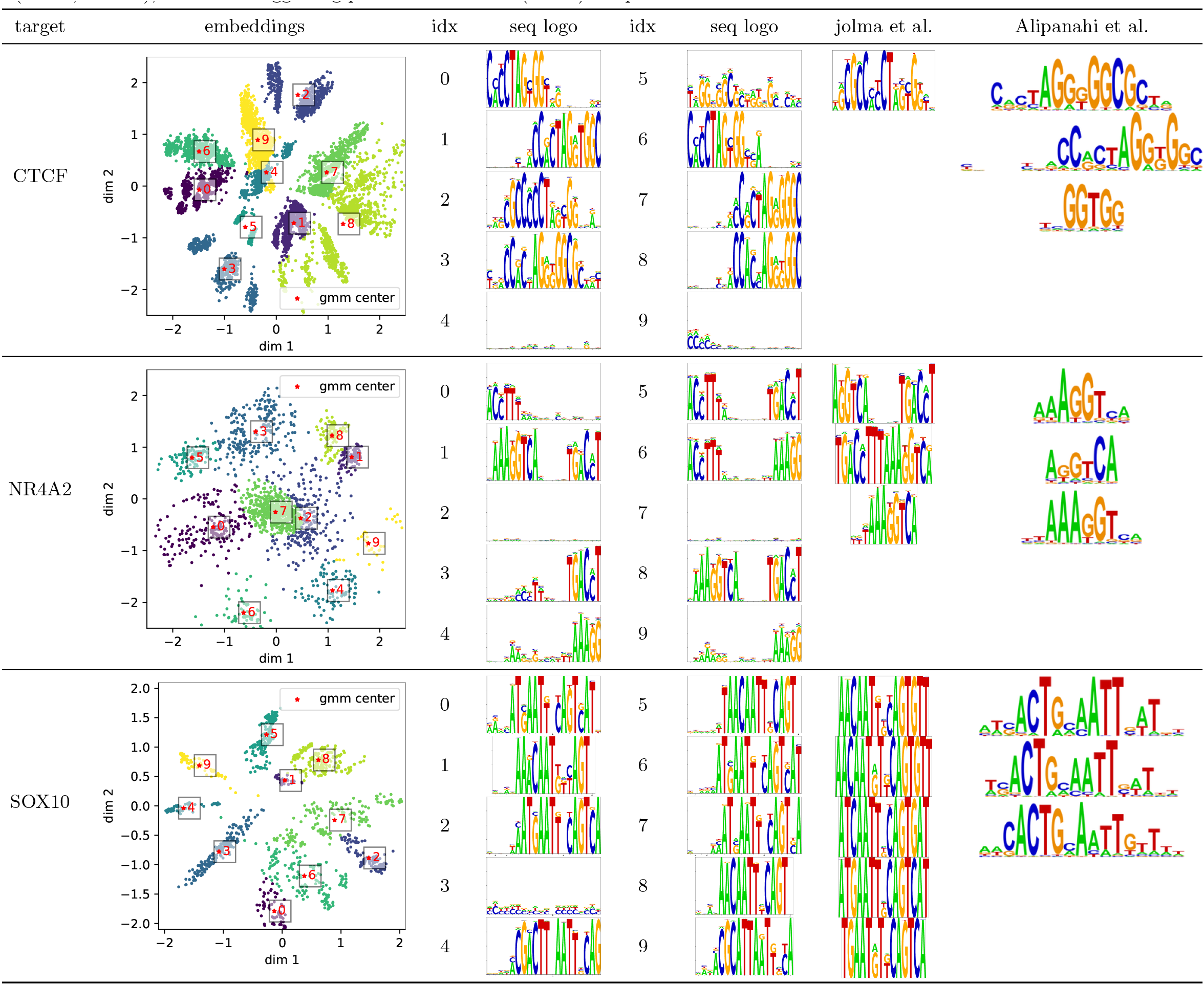
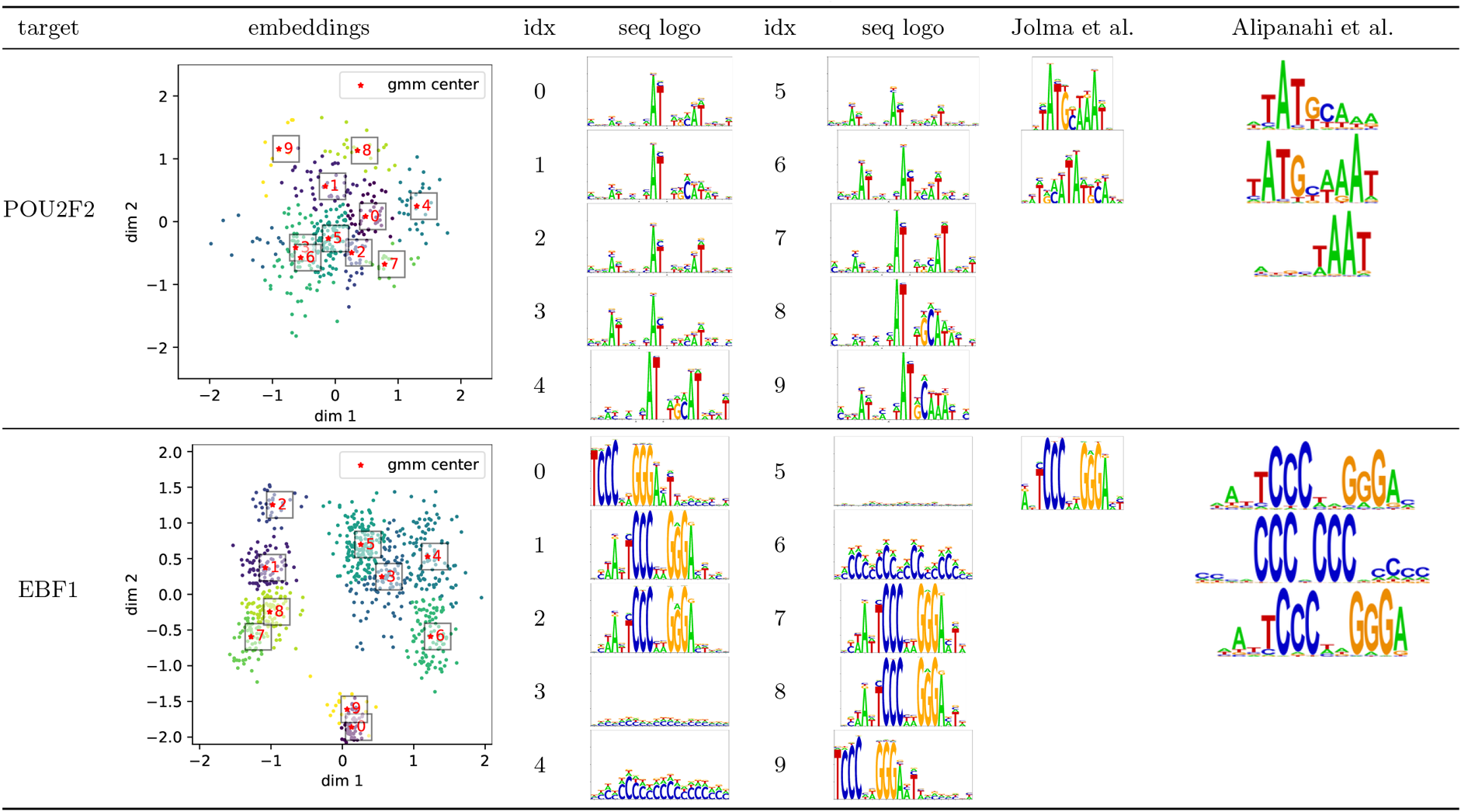
Sequences taken from the SELEX experiment in a previous study [8] are shown. Position interdependent motifs (CTCF, NR4A2), secondary motifs(Sox10, Pou2f2), and motif suggesting potential co-factors (EBF1) are presented.

| target | embeddings | idx | seq logo | idx | seq logo | jolma et al. | Alipanahi et al. |
| --- | --- | --- | --- | --- | --- | --- | --- |
| CTCF |  | 0 |  | 5 |  |  |  |
|  |  | 1 |  | 6 |  |  |  |
|  |  | 2 |  | 7 |  |  |  |
|  |  | 3 |  | 8 |  |  |  |
|  |  | 4 |  | 9 |  |  |  |
| NR4A2 |  | 0 |  | 5 |  |  |  |
|  |  | 1 |  | 6 |  |  |  |
|  |  | 2 |  | 7 |  |  |  |
|  |  | 3 |  | 8 |  |  |  |
|  |  | 4 |  | 9 |  |  |  |
| SOX10 |  | 0 |  | 5 |  |  |  |
|  |  | 1 |  | 6 |  |  |  |
|  |  | 2 |  | 7 |  |  |  |
|  |  | 3 |  | 8 |  |  |  |
|  |  | 4 |  | 9 |  |  |  |

Table S3 continued from the previous page
| target | embeddings | idx | seq logo | idx | seq logo | Jolma et al. | Alipanahi et al. |
| --- | --- | --- | --- | --- | --- | --- | --- |
| POU2F2 |  | 0 |  | 5 |  |  |  |
|  |  | 1 |  | 6 |  |  |  |
|  |  | 2 |  | 7 |  |  |  |
|  |  | 3 |  | 8 |  |  |  |
|  |  | 4 |  | 9 |  |  |  |
| EBF1 |  | 0 |  | 5 |  |  |  |
|  |  | 1 |  | 6 |  |  |  |
|  |  | 2 |  | 7 |  |  |  |
|  |  | 3 |  | 8 |  |  |  |
|  |  | 4 |  | 9 |  |  |  |

